# Nanometer condensate organization in live cells derived from partitioning measurements

**DOI:** 10.1101/2025.02.26.640428

**Authors:** Christina Dollinger, S. Thomas Hennigan, Evdokiia Potolitsyna, Abigail G. Martin, Nayara Alcantara-Contessoto, Archish Anand, Gandhar K. Datar, Jeremy D. Schmit, Joshua A. Riback

**Affiliations:** Department of Molecular and Cellular Biology, Baylor College of Medicine; Houston, TX 77030, United States of America; Medical Scientist Training Program, Baylor College of Medicine; Houston, TX 77030, United States of America; Department of Physics, Kansas State University, Manhattan, KS 66506, United States of America

## Abstract

Biomolecules self-organize into membrane-less organelles known as condensates that compartmentalize essential biochemical processes, such as ribosome biogenesis in the nucleolus^1–3^. Molecular dynamics within condensates are governed by chemical preferences and interaction networks that can imbue nanoscale structure^4–7^. Such organization is typically inferred from ensemble-averaged measurements, such as scattering and electron microscopy, which reveal molecular arrangements^8–14^. However, the complexity of cells obscures the interpretability of these techniques, limiting insight into condensate internal structure and roles in macromolecular assembly and transport. Here, we develop an approach to quantify the average microenvironment surrounding specific proteins within condensates in live cells, using thermodynamic principles to interpret the partitioning of designed protein probes. Using this approach, we find that condensates in cells, including the nucleolus, stress granule, and nuclear pore, exhibit spatial inhomogeneity, aligning with emerging views of condensates as networked fluids^5,6,15–18^. Within the nucleolus, we link spatial inhomogeneity to ribosome biogenesis, which progressively loosens the average local meshwork, facilitating transport of assembled ribosomal subunits. Within the nuclear pore, we find that transporters experience a weaker local meshwork than nucleoporins, consistent with the selective phase model^19,20^. Together, our approach uncovers a distinct mode of biomolecular control arising from nanoscale structure, which we term *microenvironment coupling*, whereby internal interaction landscapes shape transport to enable regulation and proofreading.

## Main

Organized molecular transport mitigates the stochasticity of diffusion, enabling cellular function. In the nucleus, the biogenesis of diverse RNA-protein complexes occurs via bidirectional trafficking within biomolecular condensates. As membrane-less organelles, condensates require internal cohesivity arising from solvent immiscibility and associative interactions^8^. A central question is how cohesivity gives rise to microenvironments that regulate molecular transport and biochemical function^21–23^. Limitations in resolving the organization of biomolecules within condensates hinder understanding of how condensates impact biochemical processes in physiology and disease.

Recent computational and reconstitution studies have uncovered internal heterogeneity and density fluctuations within condensates^6,8,15,16,24,25^. Given that condensates possess distinct chemical composition (e.g., ionic conditions and pH) and meshwork structure^5,17–20,23,26–32^, these findings raise the possibility of spatially heterogeneous microenvironments within a single phase of a condensate in cells. However, directly resolving and validating such microenvironments in endogenous condensates within living cells remains a central challenge.

Classical methods for resolving the structure of materials are poorly suited to study condensates in cells. Small-angle scattering and electron microscopy, among the most powerful approaches for studying liquid-like organization, cannot resolve specific proteins within cells, limiting them to highly simplified model condensates^6,9,11,12,33^. In contrast, microrheological methods, such as FRAP, fluctuation spectroscopies, and single-particle tracking, measure apparent viscosities and infer meshworks but are challenging to interpret^8,28,34,35^. Spatial proteomics methods provide information on protein abundances and localization, not nanometer structure^36^. To address these limitations, we develop strategies to elucidate the internal organization within a condensate at nanometer resolution in live cells.

## Extracting partitioning energies in live cells with microscopy

We sought to infer condensate internal structure in living cells from partitioning measurements, reasoning that partitioning provides a thermodynamic readout of the microenvironments within a condensate (**SI 4**). This approach is inspired by experiments that established general principles of protein stability, including partitioning measurements of model peptides between solvents^37–41^. However, unlike these idealized measurements, condensates in cells are highly composition-dependent^42^, requiring partitioning measurements at near-zero expression to preserve endogenous properties (**Fig. 1A**). To proceed, we initially focused on the granular component (GC) of the nucleolus as a proof-of-principle system, as the GC is a well-established condensate with a proteome and transcriptome linked to ribosome biogenesis, and with nanometer-scale features, including EM-observable substructure, that can aid validation^1,43,44^. Notably, we previously found that ribosomal intermediates diffuse ∼5,000-fold more slowly than nucleolar proteins^4^; however, the structural properties that underlie this substantial difference in transport remains unclear.

**Figure 1:**
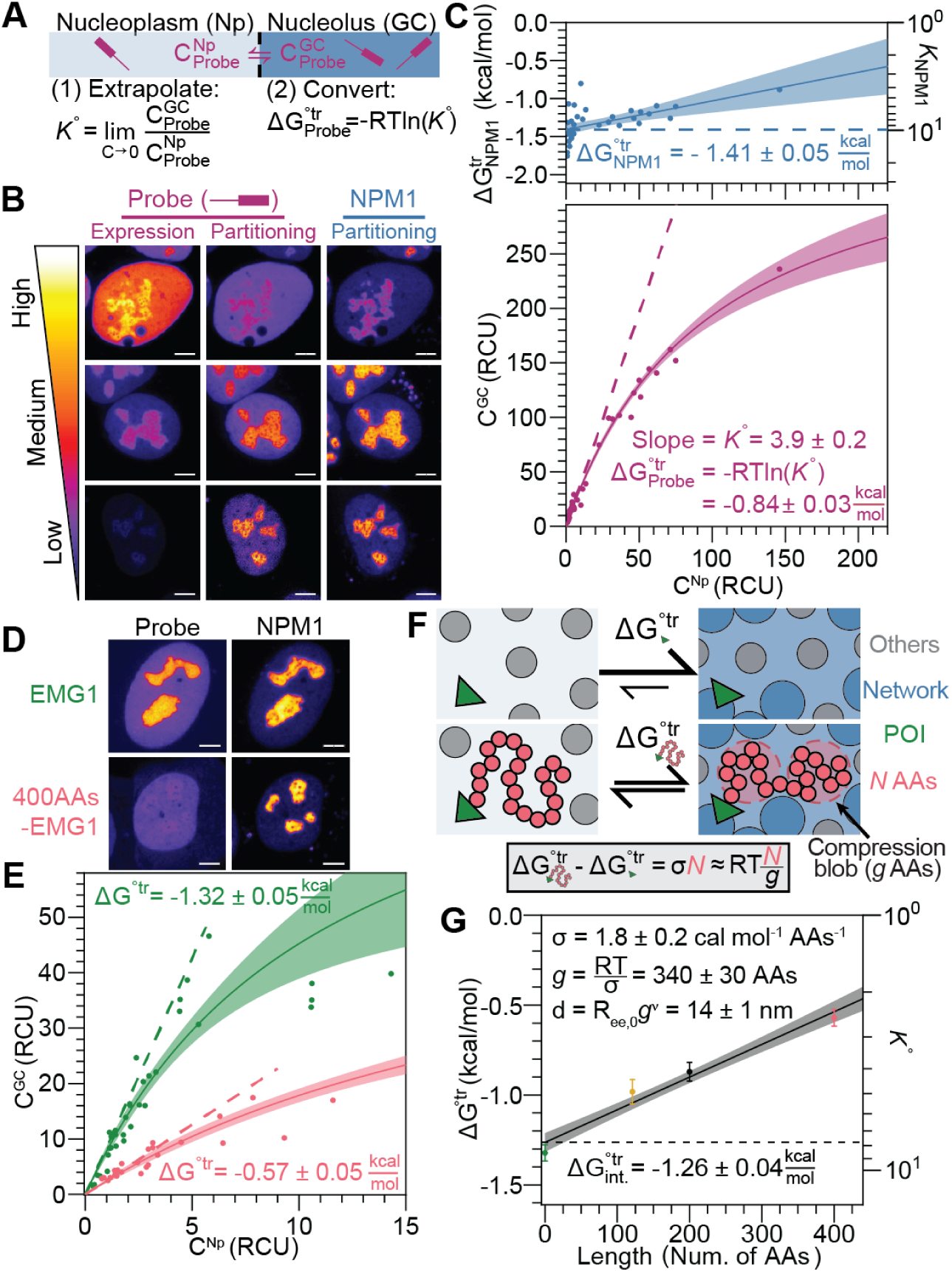
Local size exclusion measures the effective mesh size around a specific protein within a condensate. (**A**) Standard state transfer free energy measurement requires the determination of the concentration, *C*, of a probe molecule between compartments. (**B**) Various expression levels of NPM1 C-terminal truncation with mGreenLantern (mGL), the probe, in U2OS with endogenous NPM1-mCherry. Partitioning LUTs are scaled relative to the Np intensity in each image, with the upper bound set to a shared partitioning value for that channel. (**C**) Quantification of the probe levels (bottom) and the endogenous NPM1 Δ*G^tr^*(top) between the GC and Np phases. Dashed lines show trends extrapolated to zero probe overexpression. Relative concentration units (RCUs) are the number of photons emitted at reference settings (methods). (**D**) Cells with low overexpression of indicated constructs. (**E**) Quantification of indicated EMG1 probes. (**F**) Transfer free energy measurement for a protein of interest (POI) without and with an *N* residue chain. (**G**) Δ*G*^◦^*^tr^* dependence on the number of chain residues added to the EMG1 probe. Calculation of the effective mesh size shown (inset). Unless indicated, all images are fire mode, partitioning scaled, and scale bars are 5 µm.

To enable quantitative microscopy of the GC, we generated a homozygous knock-in of mCherry at the C-terminal end of NPM1, the most abundant GC protein (**Fig. S1A)**^42,45,46^. The NPM1-mCherry signal renders the nucleoplasm (Np) and GC readily identifiable, enabling quantification of concentrations (C) in each region and their ratio, the partition coefficient *K*. These NPM1-defined regions were used to quantify exogenously expressed fluorescent protein fusions, hereafter referred to as ‘probes.’ (**Fig. 1A-C**).

Having established a framework for defining compartments and measuring concentrations (methods), we next examined how probe expression impacts NPM1 and probe localization. To gain universal insights, we quantified 148 probes designed for the GC, measuring each probe in ∼30 cells over a large expression range (**Fig. S1B**). At low expression, all probes showed linear dependence, indicative of a consistent partition coefficient across cells. However, at higher expression, many exhibited a decreased slope, corresponding to a lower partition coefficient and approaching a max probe concentration in the nucleolus (**SI 2**). Consistent with a site saturation model (**SI 3**), the dependence fit well to a hyperbola, yielding two quantitative parameters: *K*^◦^, the partition coefficient extrapolated to zero probe expression, and the max probe concentration in the GC (methods, **Fig. S1B-C**). To quantify the average energetic favorability of a probe partitioning into the GC, we convert *K*^◦^ into a standard-state transfer free energy (Δ*G*^◦^*^tr^*), using the relationship, Δ*G*^◦^*^tr^* = -RTln(*K*^◦^). We denote these quantities with “◦” to indicate extrapolation to zero probe expression (**SI 1**). This extrapolation mitigates well-known consequences of overexpression^5,42,47^, including changes to endogenous NPM1 partitioning (**Fig. 1C, top**). This framework enables direct quantitative assessment of condensates without perturbing their native state.

### Measuring effective mesh size with *Local Size Exclusion* (*LSE*)

Equipped with a procedure to measure the standard-state transfer free energy (Δ*G*^◦^*^tr^*) of a probe in cells, we developed an approach to infer the average local environment surrounding a protein within a condensate by comparing the Δ*G*^◦^*^tr^* with and without the addition of a specifically designed chain. In this framework, chain addition can be cast as a Widom insertion^48^ where, in the linear response regime, the slope defines the σ-value, a thermodynamic measure of the average physicochemical environment surrounding the protein (**SI 4-5**). To test this principle, we used EMG1, an essential accessory factor of ribosome biogenesis, expressed either alone or as a C-terminal fusion with a 400-residue non-hydrophobic amino acid (AA) chain. Adding the 400AA chain reduced EMG1 enrichment in the GC with *K*^◦^ decreasing from 8.5 ± 0.6 to 2.5 ± 0.2 Δ*G*^◦^*^tr^* changing from -1.32 ± 0.05 kcal/mol to -0.57 ± 0.05 kcal/mol (**Fig. 1D-E**). This change corresponds to a σ-value of 1.8 ± 0.2 cal/mol per AA, quantifying the thermodynamic cost of introducing the chain into the average microenvironment surrounding EMG1 in the GC relative to the nucleoplasm.

Because the GC contains large ribosomal intermediates and multivalent scaffold proteins, such as NPM1^4,42,45,49^, it is expected to form a collective network or meshwork that locally confines appended chains. We therefore chose a random non-hydrophobic AA chain to minimize chemistry-specific effects, with NARDINI+^50^ identifying no pattern features (methods, **SI 6, Table S3**), allowing the resulting σ-value to be attributed primarily to *Local Size Exclusion* (*LSE*). LSE can be quantified as the free energy of confining a chain within a polymer or pore network, where the mesh size is defined by the average region encompassing a chain of *g* residues, known as the compression blob^13,14,51,52^ (**Fig. 1F**). The confinement free energy 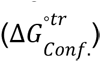 for a chain of length *N* scales with the number of compression blobs, *N/g*, yielding 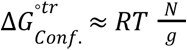; correspondingly, the LSE σ-value is 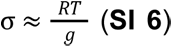. Thus, the σ-value allows us to calculate *g*, and, by employing scaling laws of unfolded proteins in water^53^, the (average) effective mesh size diameter, *d* (i.e.,d ∝ g^v^), empirical estimation suggests approximations to be near experimental error (**Fig. 1G, SI 6.1-6.3**). Furthermore, the intercept, 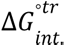, provides information about the average differences in affinity between the phases, including changes in chemical miscibility and binding interactions. To corroborate this formalism, we conducted additional measurements with two EMG1 probes: one fused with only half of the 400AA chain and another fused with the disordered region of NPM1 (121AAs), a representative disordered protein for the GC^26^. These additional measurements yield a consistent σ-value equating to a mesh size around EMG1 of 14 ± 1 nm (**Fig. 1G**). Indeed, the observation of a meshwork around EMG1 in the GC is consistent with the notion of the nucleolus as a condensate^1^.

To examine the generality of the EMG1 results as well as the heterogeneity of microenvironments in the GC, we applied LSE to nucleolar proteins, focusing on proteins with various roles in ribosome biogenesis that occur in the nucleolus^54^: (1) additional accessory factors NGDN, IMP3, and RCL1, (2) early ribosomal proteins RPL7L1 (uL30), RPS14 (uS11), and RPS11 (uS17), and (3) late ribosomal proteins RPL10A (uL1), RPS3 (uS3), and RPL12 (uL11) (**Fig. 2A, S3A**). Expression of these proteins alone (i.e., without a chain) yielded a statistically significant and expected^42,49^ correspondence between the Δ*G*^◦^*^tr^* and the ranked role of ribosome biogenesis (p<0.001, **Fig. 2B-C, S3B**). Additionally, like EMG1, adding chains to each of these proteins decreased GC partitioning (increase in Δ*G*^◦^*^tr^*) proportional to chain length, allowing for the measurement of the σ-value (**Fig. 2B, S2**). If the nucleolus was a homogenous liquid, the mesh size and, therefore, the σ-values are expected to be nearly constant across all proteins (**SI 7**)^28^. Strikingly, the σ-values (slope) ranged from 0.1 to 4 cal/mol/AA, corresponding to effective mesh sizes of 10 to 50 nm, and correlated with the ranked stage of ribosome biogenesis (p<0.01, **Fig. 2C**). The late r-Protein outlier (RPL10A/uL1, **Fig. 2C**), has recently been shown to bind rRNA at an early stage of biogenesis^55^. The 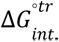 and σ-values were significantly correlated (p<0.001), consistent with a model whereby the valence of the ribosomal intermediate (inferred vis-à-vis 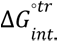) dictates the number of mesh contacts (∝ σ^3ν^) and vice-versa (fit in **Fig. 2C**, methods, also **Fig. S4**). These data are inconsistent with the GC being a homogenous liquid and instead suggest a gradual change in the local microenvironment dictated by the progression of ribosome biogenesis (**SI 7-8**). Notably, the upper range of inferred mesh sizes approaches that of ribosomal intermediates^1^, suggesting that mesh remodeling facilitates their release.

**Figure 2:**
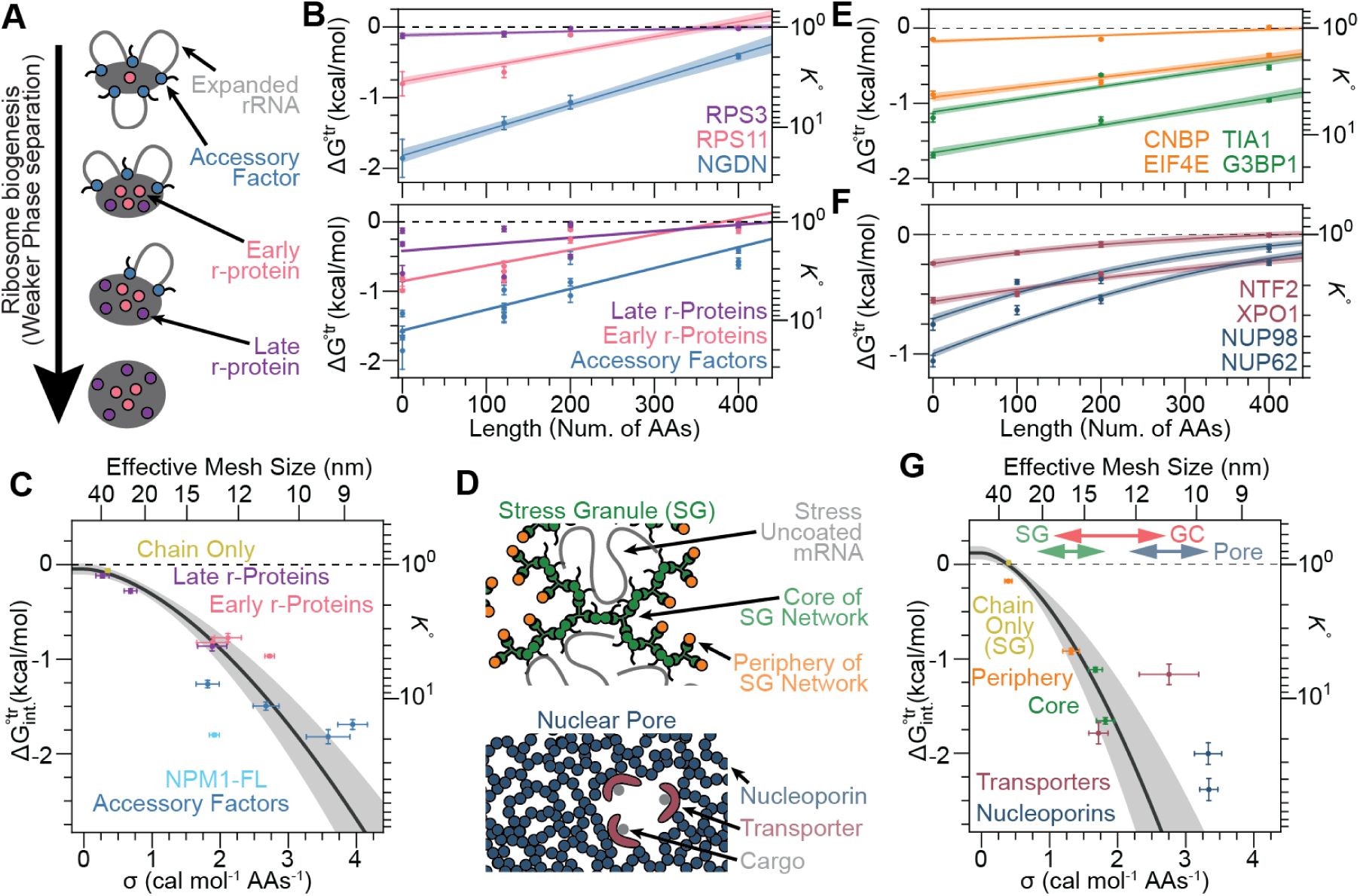
LSE reveals protein-dependent differences in effective mesh size, consistent with nanoscale heterogeneity. (**A**) Schematic of ribosome biogenesis and the categories of proteins studied. **(B)** LSE for individual (top) and categorically grouped proteins (bottom, **Fig. S3B**). (**C**) Relationship between σ and 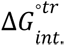 Fit assumes the 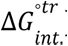 is proportional to the mesh contact number density (α σ ^3v^, methods). (**D**) Schematics of stress granules (SG) and nuclear pore (pore) networks with categories of proteins indicated. (**E-F**) LSE for individual proteins in (**E**) SG (see **Fig. S3C-D**) and (**F**) pore (see **Fig. S3E-I**). (**G**) Relationship between σ and 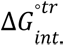 for SG and pore proteins, fit is to SG proteins. Top marks the range of effective mesh size values for indicated condensates.

We sought to further confirm the inhomogeneity within the GC. First, we focused on NPM1, a generic scaffold protein with rapid dynamics^4^ and without a direct role in ribosome biogenesis. LSE on NPM1 measured a σ-value of 1.9 ± 0.1 kcal/mol/AA or an effective mesh size of 13.2 ± 0.3 nm. These results were in line with the effective mesh sizes observed around the accessory factors and early ribosomal proteins (**Fig. 2C, light blue)**. Second, we employed the chains alone, utilizing additional random chains at each length to avoid systematic issues of charge and AA composition (**Table S3**). LSE on the chains alone measured a σ-value of 0.35 ± 0.03 kcal/mol/AA or an effective mesh size of 34 ± 2 nm in line with the late ribosomal proteins and substantially different from EMG1 and the other accessory factors (**Fig. 2C, gray)**. This further confirms the inhomogeneity in the GC.

To validate the inferred length scales and inhomogeneity, we turned to electron microscopy (EM), as the GC (“*granular* component”) is defined by EM-visible granules corresponding to ribosomal intermediates^44,56^. Although a meshwork cannot be directly resolved by EM, it should be reflected in the spatial organization of granules. Utilizing available high-resolution 4 nm isotropic EM data^57^, we identified the GC, the centers of granules, and computed the radial distribution function (RDF^4^) that measures the average signal enrichment around those centers (**Fig. S5A-E,** methods). The RDF exhibited a clear decay, followed by a well-defined minimum and a broad rise with minimal subsequent oscillations, consistent with a spatially inhomogeneous organization (**Fig. S5E**). This analysis yielded an approximate granule radius of 16 nm, in line with ribosomal intermediates^1^, along with approximate minimum and mean granule spacings of 8 nm and 26 nm, respectively, consistent with our LSE measurements (**Fig. S5F-H**). Both the inhomogeneity and inferred spacings were robust to GC identification and consistent across actively growing cell lines (**Fig. S5E-H**). Together, these orthogonal data converge on a picture of the GC as a spatially inhomogeneous network of ribosomal intermediates and support LSE as a live-cell measure of average local mesh size.

## Principles underlying LSE in endogenous condensates

Nanoscale meshwork heterogeneity may not be unique to the nucleolus, but instead represent a general feature of biomolecular condensates. We wondered whether local differences in composition and interactions create distinct microenvironments in other cellular condensates. To address this and to test the employability of LSE across condensates, we focused on stress granules and nuclear pores, by generating CRISPR-edited cell lines expressing Halo-G3BP1 or Halo-XPO1, respectively (**Fig. S3C, G**). Prior work suggests that both rely on interaction-defined meshworks; for example, FG-repeat nucleoporins form a selective, meshwork-like barrier within nuclear pores, while protein–RNA interaction networks contribute to the organization and sequestration of transcripts within stress granules^5,17–20,58,59^.

When applying LSE to stress granules, we focused on the protein interaction network that drives SG formation, including proteins at the core (G3BP1, TIA1), and periphery (CNBP and EIF4E) of this network (**Fig. 2D**)^17^ . We measured a range of mesh sizes, from ∼14 nm for core proteins to ∼40 nm for the chain alone. The σ-values for proteins at the core of the SG interaction network were higher than those at the periphery, consistent with the local meshwork being determined by network connectivity (**Fig. 2E**). As in the nucleolus, 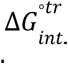 and σ-values correlated; however, fitting revealed a 2-fold steeper dependence than that observed for nucleoli, suggesting that, on average, network interactions are twice as strong to account for the larger mesh size of stress granules (**Fig. 2G**).

In the nuclear pore, we focused on transporters (NTF2 and XPO1) and nucleoporins (NUP98 and NUP62) (**Fig. 2D**). Because the nuclear pore is smaller than the diffraction limit, we globally fit the LSE data to account for the contribution of signal from the cytoplasm and nucleoplasm into that of the nuclear pore, allowing us to extract super-resolved 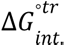 and σ-values (Fig. 2F-G, **SI 12-13**). The effective mesh size of nucleoporins was consistent with previously inferred values for passive transport through the pore, determined using dextran probes^60^. Transporters had a larger effective mesh size than nucleoporins, in agreement with the selectivity filter model^58,59^.

Similarly to the nucleolus, stress granule and nuclear pore proteins exhibited a wide variety of partition strengths 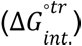 and responses to increasing chain length (σ-values), suggesting inhomogeneity (**Fig. 2E-G, SI 7**). Measured mesh sizes aligned with longstanding expectations for these condensates: stress granules were less dense than nucleoli, while nuclear pores were denser^61^. Altogether, these results demonstrate that endogenous condensates are inhomogeneous at the level of local, protein-specific microenvironments and establish LSE as a general approach to quantify them in living cells.

### Distance-dependent preferences between proteins inferred with *Length-Dependent Spatial Coupling* (*LDSC*)

Given the degree of organization implied by our LSE results, we next asked how much condensate structure dictates protein localization, focusing on NPM1. NPM1 contains three domains: an N-terminal oligomerization domain (here ‘N’) that self-pentamerizes, an IDR (‘L’), and a C-terminal RNA-binding domain (‘C’), also called the nucleolar localization sequence (RBD/NoLS, **Fig. 3A-B**). We sought to decompose the transfer free energy of NPM1 into the intrinsic domain contributions and energetic couplings between domains, analogous to established linkage analyses used in protein folding and multi-ligand binding^41,62,63^.

**Figure 3:**
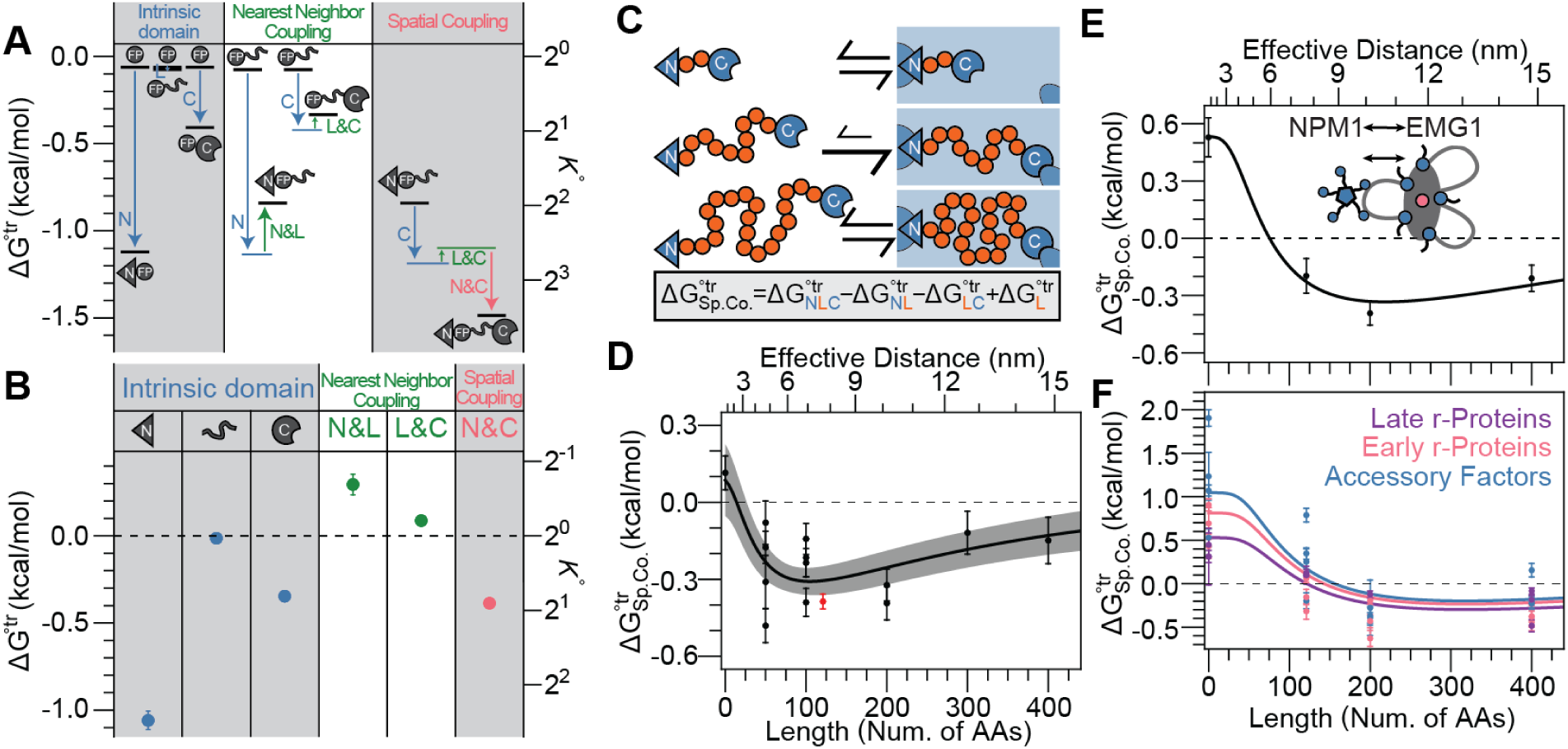
LDSC measures the effective distance between two domains or proteins. (**A**) Energetic diagram and **(B)** derived values of the domain decomposition of NPM1’s Δ*G*^◦^*^tr^*. In (A), black horizontal lines indicate measured Δ*G*^◦^*^tr^* values, and colored lines indicate theoretical values expected from indicated terms in (B). N-terminal or C-terminal addition is specified next to the arrow. (See also **Fig. S6**). (**C**) Schematic depicting the spatial coupling (Sp.Co.) as a function of chain length (top) and its calculation (bottom). (**D**) Length-dependent Sp.Co. (LDSC) between the N and C terminal domains of NPM1. NPM1’s IDR (red); not used in fit. (**E**) LDSC between EMG1 and NPM1. (**F**) LDSC between NPM1 and proteins grouped by role in ribosome biogenesis. For analytical fit form in D-F see methods.

To perform this decomposition, we measured the Δ*G*^◦^*^tr^* of NPM1-derived constructs spanning individual domains to all domains (**Fig. 3A, S6A**). Intrinsic domain contributions were determined from the difference between the Δ*G*^◦^*^tr^* of each domain and that of the fluorescent protein alone (**Fig. 3A** **left, S6A, SI 11,** methods). Nearest-neighbor couplings are the difference in the energy between a two-domain construct and that expected based on the intrinsic domain energies (**middle**). We further determined the spatial (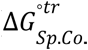, a.k.a. next-nearest neighbor) coupling which is the difference in energy between the three-domain construct and that expected based on intrinsic and nearest-neighbor domain energies (**right**).

The analysis revealed that the N-terminal domain provides the most favorable contribution to partitioning, consistent with its oligomerization with full-length NPM1. The C-terminal RBD/NoLS also contributes favorably to GC enrichment, consistent with rRNA binding (**Fig. 3B, blue-right**). In contrast, nearest-neighbor couplings weaken partitioning (**Fig. 3B, blue-middle, green, SI 9**). This is consistent with a mismatch in microenvironment preference within the GC; the termini favor local regions with smaller mesh size, which impose additional confinement on the IDR (i.e., LSE). Strikingly, the spatial coupling between the N- and C-terminal domains stabilizes GC localization, exceeding the intrinsic contribution of the RBD/NoLS alone (**Fig. 3B, magenta vs. blue-right, SI 10**). This suggests that the domains, when separated by the endogenous IDR, gain an additional energetic advantage, consistent with RNA enrichment around the pentamer. Together, these results demonstrate that partitioning cannot be explained in terms of localization sequences without consideration of local spatial organization.

Due to the strong spatial coupling between NPM1’s termini, we developed *Length-Dependent Spatial Coupling (LDSC)* to quantify distance-dependent preferences between domains (**SI 10-11**). Within this framework, spatial coupling reflects the average compatibility of microenvironments sampled by two domains as a function of their separation. Thus, coupling should depend on the average distance imposed by the intervening chain (hereafter called linkers, “L”), analogous to a distance-dependent organization function (g(r), **SI 10**). For a polymeric linker, this preference is expected to weaken when the chain is either compressed or extended away from its preferred length due to elastic constraints (**Fig. 3C**).

To test this, we measured the spatial coupling between NPM1’s N- and C-terminal domains by systematically varying the length of the intervening disordered region using polymeric linkers (same as chains from LSE, **Fig. 3D**). Fitting revealed a preferred organization at a linker length of 130 ± 13 residues, corresponding to an effective end-to-end distance of 8.0 ± 0.4 nm. Notably, this preferred length is within error of the endogenous NPM1 IDR length (121 residues), indicating that the native architecture of NPM1 is tuned to the spatial organization of its local microenvironment. This preferred organization is consistent with LSE measurements, where differences in mesh size between full-length NPM1 and its N-terminus correspond to a comparable length scale (∼5 ± 1 nm, **Fig. S3B**). Together, these results demonstrate that LDSC captures the spatial organization of microenvironments within condensates and suggest that protein architecture is matched to these nanometer-scale structures.

We next adapted LDSC to determine how ribosome biogenesis proteins are spatially organized relative to NPM1. We designed probes containing both NPM1 and proteins of interest with various lengths of linker between them, allowing measurement of spatial coupling as a function of separation. Beginning with EMG1, spatial coupling with NPM1 started repulsive but became more favorable with increasing linker length (**Fig. 3E**). Across ribosome biogenesis proteins, spatial couplings were unfavorable without an intervening linker, with the extent of repulsion decreasing with the ranked stage of ribosome biogenesis (p < 0.001, **Fig. 3F, S6B–C,** 1.1, 0.8, and 0.3 ± 0.1 kcal/mol for the three protein types). Coupling remained weakly repulsive for linkers with an effective distance of ∼7.5 nm (0.13 ± 0.02 kcal/mol), whereas at ∼10 nm and ∼15 nm, it became favorable (-0.26 ± 0.02 and -0.22 ± 0.02 kcal/mol, respectively). Fitting globally yielded a preferential effective distance at a linker length of 340 ± 30 residues, corresponding to an effective end-to-end distance of 14 ± 1 nm (**Fig. 3F**). Consistently, coupling with the NPM1 N-terminus was repulsive for all proteins, indicating that preferred organization occurs away from the center of the NPM1 pentamer (**Fig. S6**). These results indicate that NPM1 does not directly solvate accessory factors or ribosomal proteins, in line with models in which NPM1 drives GC formation through heterotypic interactions with rRNA rather than proteins^4,42^. The increased exclusion of NPM1 during early stages of ribosome biogenesis further suggests that interactions driving phase separation are spatially separated from sites of assembly (**Fig. 3E-F**), providing a mechanism to minimize competition between phase separation and ribosome biogenesis. Together, these results demonstrate that LDSC resolves nanometer-scale spatial organization within condensates revealing how biomolecules are positioned in space.

## Conclusions

Condensates are commonly proposed to regulate biochemical reactions through enrichment and exclusion of specific biomolecules, a mode of regulation largely indistinguishable from that of membrane-bound organelles. However, this view implicitly treats partitioning as an inherent, fixed property of a molecule determined by bulk properties of the condensate. In contrast, we uncover a distinct mode of biomolecular control arising from nanometer-scale structure, which we term *microenvironment coupling*. Specifically, biomolecules are statistically coupled to distinct microenvironments within a condensate (**Fig. 4C**). We find that proteins that strengthen local interaction networks generate tighter meshworks, whereas those that do not experience looser ones. Indeed, within the nucleolus, stress granule, and the nuclear pore, the strength of partitioning is strongly correlated with the local meshwork, as quantified by LSE σ-values (**Fig. 2, SI 5-8**). Extending to multi-domain proteins and protein–protein interactions revealed that partitioning is non-additive and distance-dependent, reflecting the compatibility of local microenvironments (**Fig. 3, SI 9-11**). Thus, partitioning emerges from the collective arrangement of interacting components, consistent with predictions from field-theoretic approaches^6,15^. In this way, microenvironment coupling arises from the cohesive nature of condensates and introduces an additional structural length scale that enables cooperativity, allostery, and other emergent properties to be tuned for biological control.

**Figure 4:**
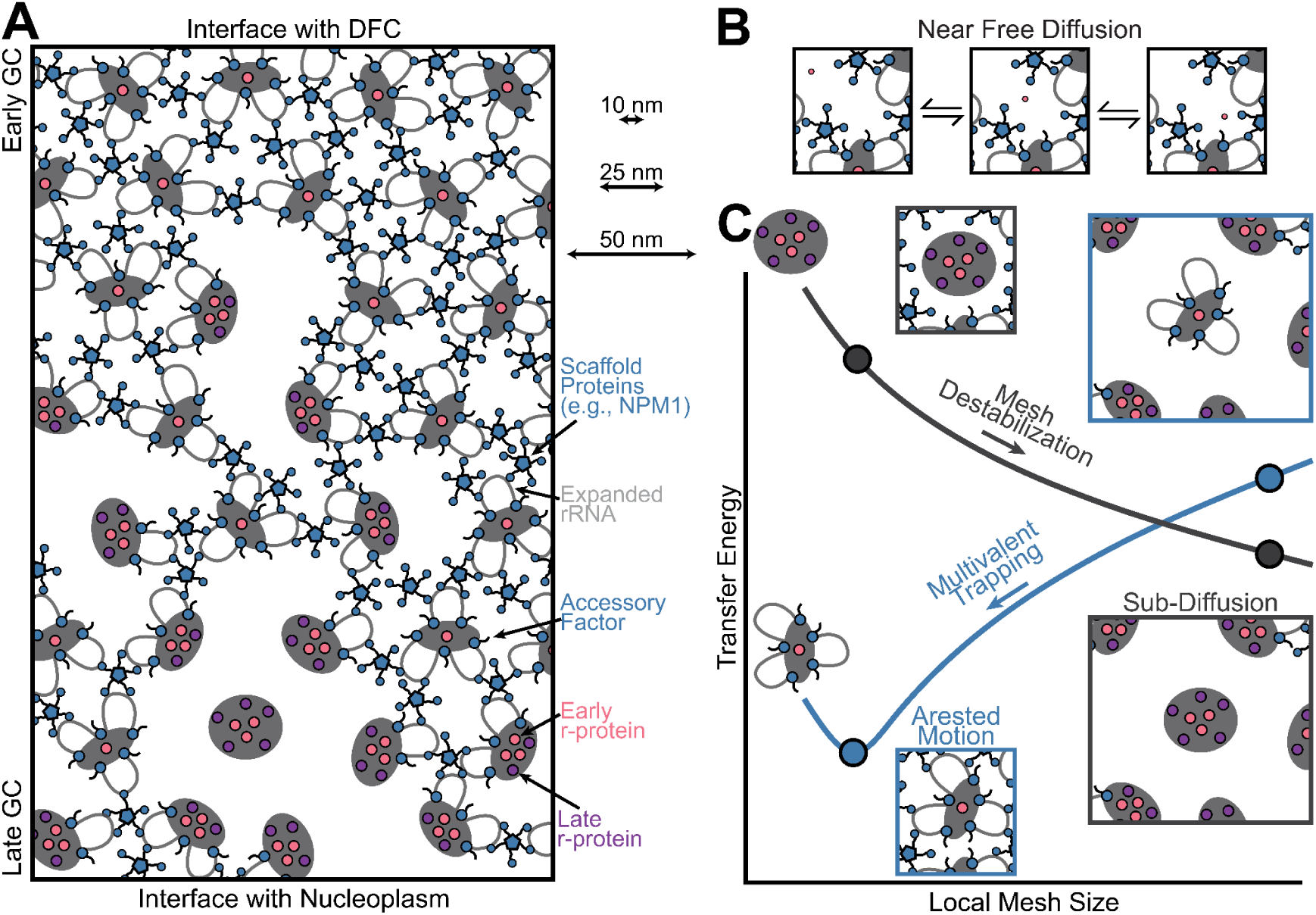
LSE and LDSC provide a nanometer picture of molecular interactions and transport during ribosome biogenesis. (**A**) Gradient of ribosome biogenesis and mesh sizes within the GC. Note that client molecules, including scaffold proteins that are in client-like states, are omitted to highlight mesh. (**B**) Depiction of free early r-protein transport within the GC meshwork. (**C**) Local mesh size dependence on transfer energy and transport for an early ribosomal intermediate (blue) and the completed ribosome (gray) due to local changes in the number of interactions (**Fig. S4**, methods).

Our measurements uncover how microenvironment coupling emerges in the nucleolus and its relationship to ribosome biogenesis (**Fig. 4**). Early GC ribosomal intermediates have high valency for phase separation, generating a more stable, tighter meshwork with a mesh size between 10-15 nm (**Fig. 2B-C, 4A**), which lies between that of a folded protein domain (∼3-5 nm) and the diameter of ribosomal intermediates (∼30 nm^1^, **Fig. 2C**). This implies that, in contrast to ribosomal intermediates, proteins can readily transport through the mesh via free diffusion or reptation for single and multidomain proteins, respectively (**Fig. 4B**), consistent with existing data^4,64^. Ribosomal intermediates initially form a structured core surrounded by multiple expanded rRNA stretches that progressively collapse during biogenesis in the nucleolus (**Fig. 2A, 4A**)^49^; we find that this step-wise process results in the sequential destabilization of the local meshwork (**Fig. S4, 4C**), eventually shifting the average local microenvironment to an inferred mesh size of ∼40-50 nm, enabling selective export of mature subunits (**Fig. 4C**). These structural inferences are consistent with the disconnect between the dynamics of proteins and ribosomal intermediates in the GC^4^, the existence of constrained (rRNA) territories around individual rDNA-containing chromosomes^65^, and the inhomogeneity of GC organization observed here and in previous EM studies^66^. Altogether, these inferences yield a picture in which microenvironment coupling endows the nucleolus with an intrinsic proofreading capacity, licensing export upon maturation.

Beyond the nucleolus, microenvironment coupling provides a general mechanism for selective transport. In the nuclear pore, the selective phase model^67,68^ can be recast within this framework, whereby transporters locally deform the meshwork, as measured here (**Fig. 2D,F,G**), enabling ratchet-like transport. We observe similar microenvironment inhomogeneity within stress granules (**Fig. 2D,E,G**) which may facilitate the reformation of translation-competent mRNPs and regulate stress granule dissolution. Together, these results imply that local and global microenvironment properties are attuned to the distinct functional roles of different condensates. More broadly, this suggests a new paradigm in which condensates control cellular biochemistry through the nanometer-scale organization of local microenvironments, with implications for disease states in which condensate organization is compromised^1,69^.

## Acknowledgments

We thank members of the Riback Lab for discussions and feedback on the manuscript. We thank Steven Boeynaems, Kyle Eagen, Alex Holehouse, Paulo Onuchic, and Tobin Sosnick for feedback on the manuscript. We thank Margaret Goodell for assistance and advice with CRISPR editing, including access to equipment. J.A.R. is a CPRIT Scholar in Cancer Research. This work was supported by funding from the Cancer Prevention and Research Institute of Texas (CPRIT) New Investigator Grant RR210040 (J.A.R.), the Ted Nash Long Life Foundation (J.A.R.), the Leukemia Research Foundation (J.A.R.), the Searle Scholars Program (J.A.R.), Welsh Foundation Q-2254-20250403 (J.A.R.), NIH F30CA268725 (G.K.D.), NIH CA183252 (Goodell PI), NIH R01GM141235 (J.D.S), and NIH R35GM162528 (J.A.R.). This work was also supported by the Cytometry and Cell Sorting Core with funding from CPRIT and the NIH (RP180672; CA125123, RR024574).

## Author Contributions

Conceptualization, J.A.R.; Investigation, C.D., S.T.H., E.P., A.M., N.A. A.A., and G.D.; Methodology, C.D., S.T.H., E.P., A.M., N.A., A.A., and G.D.; Data curation, C.D., S.T.H., and J.A.R.; Formal analysis and Theory, J.D.S. and J.A.R.; Writing - original draft, C.D., S.T.H., E.P., and J.A.R.; Writing - review & editing: C.D., S.T.H., E.P., A.M., A.A., G.D., J.D.S., and J.A.R.; Supervision, J.A.R.

## Data and materials availability

All datapoints can be found at the lab’s github (https://github.com/CellularPhysicalChemistryGroup/Manuscripts).

## Materials and Methods

### Cell culture

U2OS were cultured in sterile filtered McCoy’s 5A Modified Medium (1X) with L-Glutamine (Gibco 16600-082) and supplemented with 10% FBS (Avantor 89510-186) and 1% Penicillin–Streptomycin (Gibco 15140-122). HEK293 and HeLa cells were cultured with DMEM (1X) with 4.5 g/L D-Glucose and L-GLutamine (Gibco 11965-092). Cells were kept in a humidified incubator (VWR, PHCbi MCO-170AICUVDL-PA) with 5% CO_2_ and at 37°C. CO_2_ levels were calibrated every six month sterilization cycle. Temperature was checked every week using a thermometer. For routine passaging, cells were washed with PBS pH7.2 (1X) (Gibco 20012-043), dissociated with trypsin (Trypsin-EDTA 0.25%, Gibco 25200072) then plated into a T25 cm^2^ flask. For imaging, cells were seeded into an 8-well (Ibidi 80807) or 96-well plate (Cellvis, P96-1.5H-N) coated with fibronectin (VWR 103701-196), according to the manufacturer’s instructions. Mycoplasma testing on all cell cultures was conducted every two weeks using ABM’s Mycoplasma Test Kit and protocol (Applied Biological Materials G238).

### CRISPR protocol

U2OS NPM1-mCherry was originally generated in^70^. In brief, the DNA template encoding fluorescent protein mCherry was purchased from Twist Biosciences (**Table S1**). The template was PCR amplified with KAPA HiFi HotStart ReadyMix (Roche) and custom oligomer primers to add flanking 100-200bp homology arms, generating locus-specific homologous recombination repair templates (**Table S1**). Cells were electroporated with the Cas9-gRNA ribonucleoprotein (RNP) complex and the HDR template using the Neon Transfection System (Thermo Fisher) at the following parameters: 1350 V, 35 ms, 1 pulse.

U2OS Halo-XPO1 and U2OS Halo-G3BP1 were generated as follows. HDR templates were amplified from plasmids containing selected fluorophores using Roche KAPA 2x DNA Polymerase (**Table S1**). HDR templates were then purified using the AMPure XP beads with size selection for longer fragments. For editing, 250,000 cells per reaction were mixed with RNP complexes (2 ug Cas9, 2 ug gRNA), 2 ug HDR fragments, and electroporated using the Lonza 4D-Nucleofector with the SE kit and CM-104 program. Following electroporation, cells were expanded, and positive cells were enriched by fluorescence-activated cell sorting using a Sony MA900 cell sorter in an isotonic buffer containing 5% FBS. A minimum of 3000 cells was used to establish stable cell lines, which were further validated by imaging and PCR genotyping across the insertion site. Genomic DNA was extracted using a NEB kit (T3010L), and target loci were amplified via PCR using genotyping primers (**Table S1**). The PCR products were then analysed by agarose gel electrophoresis (**Fig. S1A**).

### Probe generation

#### Selection of proteins of interest

The ribosome biogenesis proteins employed were identified for their experimentally determined roles in this process (accessory factors, early, and late ribosomal proteins)^54^. For stress granules, we focus on proteins spanning the protein-protein interaction network^17^, selecting G3BP1 and TIA1 from the core of the network and CNBP and EIF4E from its periphery, the latter being associated in 5’ TOP mRNA translational regulation^71^. For the nuclear pore, we selected the FG-repeating nucleoporins, NUP98 and NUP62, and transport factors, XPO1 and NTF2, that interact with and traverse through the barrier. The fluorophore and chain tag sites were chosen based on previous literature or on the OpenCell database (**Table S4**). Sequences of canonical isoforms were amplified from the cDNA of U2OS and HEK293 cells and used as inserts for further cloning into an overexpression vector.

#### Fluorophore selection

To ensure that the fluorescent protein on the probes did not bias apparent partitioning by causing enrichment or exclusion, we screened fluorescent tags expressed alone in condensate-marked U2OS cells to identify those with a K≈1. mGreenLantern and mStayGold (later used for expanded LSE measurements in stress granules and at the nuclear pore), both showed partition coefficients near 1, indicating minimal enrichment or exclusion from the condensates examined (**Fig. S2**).

#### Chain design

Four 100-residue chains (L100A-D) were randomly generated using Mathematica with an equal probability of all amino acids except for aliphatics and aromatics (alanine, valine, leucine, isoleucine, methionine, cysteine, phenylalanine, tyrosine, tryptophan, and histidine) (**Table S3**, see code). DNA encoding the four sequences were ordered from Twist Bioscience and cloned into the vector at the chain insertion site. After sequence verification, chain-alone constructs were transfected into U2OS cells to validate expression and confirm the absence of substantial enrichment or depletion in the nucleolus, stress granules, and nuclear pore. The longer chains used throughout this paper were assembled from these four (**Table S3**), and similarly tested to ensure no enrichment or depletion. To examine whether the chains inadvertently encoded endogenous IDR-like patterning, we analyzed them with NARDINI+^50^, which assesses whether residue classes are more well-mixed (indicated by negative Z-scores) or more blocky (positive Z-scores) than expected from same-composition sequence scrambles. The L100A-D chains lacked strong pairwise patterning signatures: L100A and L100D had no absolute value Z-scores above 1 (indicating a random sequence), and L100B and L100C scored only modestly above the threshold (<1.1), except that L100B showed a 1.7 in a positive-positive residue pattern. When examining NPM1’s length dependence partitioning in LDSC (**Fig. 3D**), we used all 100-AAs chains, and when designing probes for LSE on the nuclear pore (**Fig. 2G**), we opted for L100A because of its low patterning feature Z-scores. Together, the sequence analysis and chain-alone localization controls indicate that the chains were broadly grammar-benign for the intended length-control measurements.

#### Construct cloning

We engineered a custom lentiviral cloning vector, pTwist Lenti SSFV, in which the native multiple cloning site was replaced with a modular cassette containing three independent Gibson assembly insertion sites (termed N-terminal (NheI), Chain (AfeI), and C-terminal (SpeI)) around a fluorophore; **Table S2**). Each site was designed with unique, non-overlapping homology regions (**Table S5**), allowing inserts to be introduced at any position without affecting the remaining sites. This design enables sequential or combinatorial assembly of multiple elements within the same backbone. A monomeric fluorescent protein mGreenLantern^72^ or mStayGold^73^ was positioned immediately upstream of the Chain site, allowing fusion constructs to be generated without additional subcloning.

Vectors were linearized using NheI (NEB R3131L), AfeI (NEB R3552S), or SpeI (NEB R3133S), depending on the target insertion site, and assembled with PCR-generated inserts carrying the appropriate homology arms using Gibson Assembly Master Mix (NEB E2621S). The plasmid constructs were transformed into NEB Stable Competent E. coli (High Efficiency) (C3040H), following the manufacturer’s guidelines, plated onto Agar plates with Ampicillin (Thermo-Fisher Scientific 11593027), and left overnight in a 37 C° incubator. The following morning, single colonies were isolated, expanded in LB media and Ampicillin overnight at 37 C° shaking, and mini-prepped with Monarch Plasmid Miniprep Kit (catalog T1010). All constructs were verified by full-plasmid sequencing (Plasmidsaurus). In practice, using the same plasmid backbone and restriction digestion sites across all probes allowed us to rapidly generate the 186 constructs used in this analysis.

#### Plasmid Preparation

Once sequenced-verified, all plasmid constructs were purified using the ZymoPURE Express Plasmid Midiprep Kit (Zymo Research, D4213-A) following the manufacturer’s instructions. Bacterial cultures were grown overnight in LB medium containing Ampicillin to induce growth, and plasmid DNA was eluted with sterile water and quantified by absorbance at 260 nm prior to transfection.

### Transient Transfection of construct probes

Cells were seeded at 20-25,000 per cm^2^ in an 8-well or 96-well plate. The next day, following the vendor’s vendor’s standard protocol, cells were transfected with 0.25-0.5µg plasmids (midi-prep concentrations at 500-5000 ng/µL), 1 µL Lipofectomine P3000 reagent, 0.75 µL Lipofectomine (Thermofisher L3000008), and 25 µL Optimem 1X (Gibco 31985-070) per well. Following 5-hour incubation at 37 °C, transfection media was replaced with complete growth media (McCoy’s). Cells were imaged 48-72 hours after transfection.

### HaloTag–JF549-HTL Labeling

U2OS cells expressing HaloTag-fusion to either XPO1 or G3BP1 were labeled with the cell-permeable fluorescent ligand JF549 HaloTag dye (Promega, HT1020). Cells were incubated in a complete growth medium containing 100 nM JF_549-HTL_ for 30 min at 37 °C, protected from light. Following incubation, cells were washed three times with pre-warmed PBS to remove unbound dye and then allowed to recover in dye-free growth medium for 15–30 min at 37 °C before imaging. This washout period ensured complete removal of nonspecific fluorescence and allowed equilibration of labeled proteins.

### Stress Granule Induction

After labeling HaloTag-G3BP1 in U2OS cells, stress granules were induced with sodium arsenite (Millipore-Sigma, S7400) at a final concentration of 0.5 mM in growth medium at 37 °C. This acute oxidative stress treatment reliably triggers the formation of cytoplasmic stress granules, as confirmed by the redistribution of our HaloTag-G3BP1 marker. Cells were imaged and analyzed 1 hour after induction.

### High-resolution live-cell microscopy

All images were taken on a microscope setup containing a Nikon’s Ti2E, a VisiTech instant SIM microscope (VT-iSIM), a Hamamatsu ORCA Quest qCMOS camera, and a CFI60 Plan Apochromat Lambda D 100X oil immersion objective lens. Cells were maintained in a 5% CO_2_ at 37°C chamber and imaged with immersion Oil Type B 37°C (Nikon 77005). Imaging of green (mGreenLantern) and red (NPM1-mCherry) was done sequentially via excitation with 505 nm and 561 nm laser, respectively. No detectable bleed-through was observed. For stress granules and the nuclear pore, the endogenous markers (G3BP1 and XPO1, respectively) were fused to HaloTag, and were imaged at 488nm (mStayGold) and 561nm (JF_549-HTL_), sequentially.

Power meter measurements were taken at the focal plane for both channels before imaging to verify the proper alignment and performance (e.g., sufficient warming up). Dye standards were also used approximately once a month to determine the no-emission background (due to factors such as scattering of the excitation light) and to measure the extent of vignetting needed for flat field correction for all lasers at all power settings. Additionally, this procedure determined the digital level (DL) offset and the conversion between DL and photoelectrons (via fitting low light photon counts histogram) for all camera settings. The camera linearity was verified manually by confirming that pixel values increased proportionally with increasing laser power, indicating that the detector was operating within its linear range and below saturation.

### Determination of ROIs and calculation of RCU

All images were analyzed using a custom plugin in Micro-Manager and ImageJ/Fiji. Regions of interest (ROIs) corresponding to dense and dilute phases were defined using the endogenous compartment markers (NPM1-mCherry for nucleolus, Halo-G3BP1 for stress granules, and Halo-XPO1 for nuclear pores). Dilute-phase ROIs (nucleoplasm or cytoplasm) were selected manually as polygons or circles, avoiding out-of-plane structures. Dense-phase ROIs were either a 5-pixel radius gaussian ROI or square ROI, algorithmically chosen as the ROI with the maximum compartment marker signal within the cell. Square ROIs were 5x5-pixel sizes for nucleoli and stress granules but reduced to 3x3-pixel sizes for the nuclear pore, lowering the size due to the diffraction-limited nature of the pore. All automated selections were manually validated to ensure correct compartment assignment, with search regions adjusted in the rare cases selection was incorrect (e.g., not within the cell). ROI selection was performed blinded to the probe channel wherever possible. Relative concentration units (RCUs) were then determined by averaging the DLs within the ROIs, subtracting the no-emission/no-sample background, and dividing by the average flat-field calibration. Subsequently, a sample-specific background was also subtracted. For each cell, a single RCU value was obtained per compartment. Measurements were reproducible across experimental days, including comparisons spanning over one year.

### Extraction of the steady-state transfer free energy

For each construct (i.e., probe), dense- and dilute-phase concentrations (RCUs) were obtained from ∼30–50 cells spanning a broad range of expression levels (12.2, 577.5, and 59.2 fold for 2, 98, and 50 percentiles). To extract the steady-state partition coefficient, we modeled the dense-phase concentration as a function of dilute-phase concentration using a two-parameter saturation model (rectangular hyperbola; **SI 2-3**). This yields the steady-state partition coefficient (*K*^◦^), which corresponds to the fit slope extrapolated zero expression levels, and can be employed to calculate the transfer free energy (ΔG°^tr^). Note, inclusion of low-expression cells is essential to ensure robust extrapolation. While capturing the full (hyperbola) dependence is ideal, probes with near-unity partitioning (i.e., minimal enrichment or exclusion) often remain effectively linear even at the highest expression levels achievable with our promoter (50% of constructs, **Fig. S1**).

Fitting was implemented in Mathematica using an iterative nonlinear weighted regression routine. Initial parameter estimates were obtained from a filtered, smoothed version of the dataset, using a moving-median procedure to reduce the influence of outliers. The model was then refined by iterative reweighting on the full dataset. At each iteration, residual error was quantified as the approximate squared orthogonal distance from each data point to the fitted curve, rather than as a purely vertical deviation to account for uncertainty in both variables, and the median of these errors is used to set the characteristic error scale. Points with large deviations from the current fit were progressively down-weighted, as they typically represent out of plane dense-phase ROIs. Low-expression measurements are additionally up-weighted (with weight inversely proportional to the dilute concentration) to ensure that the extrapolated low-expression slope is constrained by the data most relevant for extracting *K*^◦^. The nonlinear optimization was performed with a Levenberg–Marquardt routine until convergence. In a small number of cases (<10), manual constraints on the fitting range and/or model form were applied. These instances were identified by visual inspection, where the automated iterative routine did not adequately capture the low-expression regime, typically due to sparse sampling or disproportionate influence of higher-expression data. In such cases, fitting bounds or linear constraints were imposed to ensure that the low-expression behavior was accurately represented. See code for exact Mathematica functions.

### Extraction of sigma-value

For each protein, σ-values were determined by measuring the steady-state transfer free energy (ΔG°^tr^) of the probe with appended disordered chains of varying length, as described above. Briefly, this approach measures ΔG°^tr^ for a given protein as a function of the length of an added chain, where the dependence on chain length captures the local microenvironment surrounding the protein (**SI 5-7**). At minimum, two chain lengths are required to extract σ and 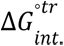; practice, additional intermediate chain lengths were included to ensure robust determination. In most cases, four chain lengths were used. For each protein, ΔG°^tr^ values from all corresponding chain variants were compiled and fit as a function of chain length, with σ defined as the slope and 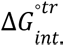 as the intercept.

This fit was implemented in *Mathematica* using linear regression. All chain variants were included, with each construct contributing a single data point, and ΔG°^tr^ uncertainties propagated from the fits described above. A robust variance estimator, defined as the median squared uncertainty within each set, was used, and confidence intervals were calculated at the 68% level. The σ-value was taken as the fitted slope, with uncertainty obtained from the regression. The intercept was extracted from the same fit. σ-values were subsequently converted into effective mesh sizes as described in the main text and **SI 6**. For comparison of the linear with the non-linear dependence of confinement, we fit the interpolated Casassa equation similarly (**SI 6.2**). See code for exact Mathematica functions..

### Relationships between sigma, local mesh contact density, and the transfer free energy

To understand the relationship between the σ-value and the transfer free energy of a protein (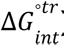), we took a mean-field approach. In this framework, 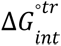 is proportional to the average local mesh contact density (δ). Using scaling arguments, the contact density is expected to scale as the inverse cube of the mesh size, δ ∝ *d*^−3^ . The mesh size is related to g and σ, *d* ∝ *g^v^* ∝ σ^-v^, where ν is the Flory exponent for a protein in water, approximated as 0.55 (**SI2.6**). Together, this yields δ ∝ σ^3v^, which relates the σ-value to the local mesh contact density. We therefore fit 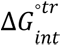 as linear function of σ^3v^ minimizing the sum-of-squares error with 50 iterations to propagate errors using the best fit from previous iterations. After including necessary conversions, σ can be mapped to the local mesh contact density (δ) via the relation δ = 0. 25 σ^3v^ having units of millimolar, yielding values of 1.5, 1, and 0.2 mM for the accessory factors, early r-Proteins, and late r-Proteins respectively. The resulting slope for nucleolar proteins is 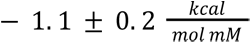, while stress granules exhibit a steeper dependence 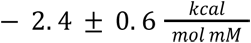, indicating stronger effective interactions per unit contact density.

Extraction of spatial coupling from linear decomposition of transfer free energy by domains.

To determine the spatial coupling at a fixed linker length between two domains of one protein or two different proteins, four constructs are required, derived from the three domains, N-terminal domain/protein, linker, and C-terminal domain/protein abbreviated N-L-C for the full construct. The four measurements are 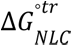 on the full length construct, 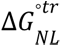 and 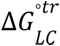 on the two-domain constructs containing the linker, and 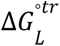 on the linker alone. The spatial coupling calculation from these four constructs is 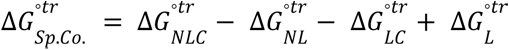. Because the resulting error is determined by sum of squares 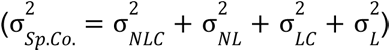, it is essential that the error on each construct be substantially low. A full description of the theoretical description of the spatial coupling and its derivation are provided in **SI 10-11**.

The intrinsic domain transfer free energy, 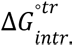, is calculated as the difference between the transfer free energy of the domain (with a fluorescent protein) and that of the fluorescent protein alone. Similarly, the nearest-neighbor coupling transfer free energy, 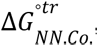, is calculated as the difference between the sum of the transfer free energies of the two-domain construct and that of the fluorescent protein from the sum of the transfer free energies of the constructs containing the individual domains. For example for the nearest-neighbor coupling between the N-terminus and the linker, the equation is 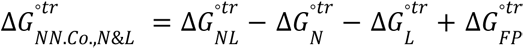. In each case the errors were calculated by the sum of squares. A full description and derivation can be found in SI 11.

### Fitting the LDSC

To quantify distance-dependent interactions between domains or proteins, we model spatial coupling as arising from the convolution of polymer conformational statistics with a distance-dependent interaction function. Specifically, the spatial coupling free energy is expressed as an integral over distances weighted by the end-to-end distribution of the intervening chain, P(r|N), and a radial interaction function analogous to a pair-correlation function. See **SI 9-10** for more information. In practice, we approximate this behavior using a simplified model consisting of two contributions: (i) a distance-independent term capturing interactions that do not depend on domain separation, and (ii) a distance-dependent term describing a preferred interaction at a characteristic separation. The latter is modeled assuming a Gaussian chain distribution^13,14^, yielding a functional dependence on chain length N with a minimum at a preferred length scale. Thus:

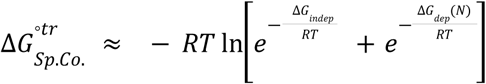

with

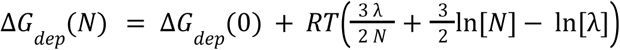

where λ is the length of AAs at the preferred distance and *N* is the length of the chain. Additionally, to account for a few chain residues of linker and the fluorescent protein, the equation is fit with *N* values increased by 25 residues. When globally fitting for NPM1 LDSC we add a σ^3ν^ dependent term to Δ*G_indep_* (**Fig. S6C**).

### Electron Microscopy analysis

Electron microscopy data were analyzed from publicly available volume EM datasets of interphase HeLa and Jurkat cells^57^. Single 2D slices containing nucleoli with clearly identifiable granular component (GC) ultrastructure and adjacent nucleoplasm were extracted from the 3D datasets and exported as CSV files. All analyses were performed in Mathematica (link found in Data and materials availability). See code for exact implementation, including functions emImportData, calcCenters, figureRDF, and getLocation, which perform data import, granule center detection, RDF calculation, and extraction of characteristic length scales, respectively. Briefly, for each dataset, a GC-containing region of interest was selected manually, as shown in **Fig. S5A-D**. The cropped EM image was background-offset by subtracting the minimum value of a 5-pixel blurred version of the crop. Granule centers were identified by blurring the image by 2 pixels, thresholding at the midpoint between the 0.5th and 99.5th percentile intensities, dilating the resulting binary mask with a disk of radius 2 pixels, computing a distance transform, blurring the distance map by 2 pixels, and identifying local maxima using MaxDetect. To quantify spatial organization, we computed a radial distribution function (RDF)-like profile centered on the detected granules, following a related approach^4^. Discrete annular convolution kernels were generated in 4 nm radial increments, and for each radius the mean EM intensity surrounding the detected centers was calculated and normalized by the mean image intensity of the crop, yielding the average relative EM signal as a function of distance from granule centers. The RDF was smoothed by fitting the first 189 radial positions with a Bernstein-basis spline using 29 basis functions, and confidence intervals were obtained from the fitted covariance matrix. From the smoothed RDF, we extracted three characteristic positions: the first crossing below 1, the first minimum, and the subsequent crossing back to 1. The first crossing was defined as the granule radius, the difference between the first minimum and this radius as the minimum granule separation, and the difference between the second crossing and this radius as the mean granule separation. These values were measured across multiple HeLa and Jurkat regions and averaged. This analysis follows previously published methods⁴.

## Supplementary Figures

**Figure S1:**
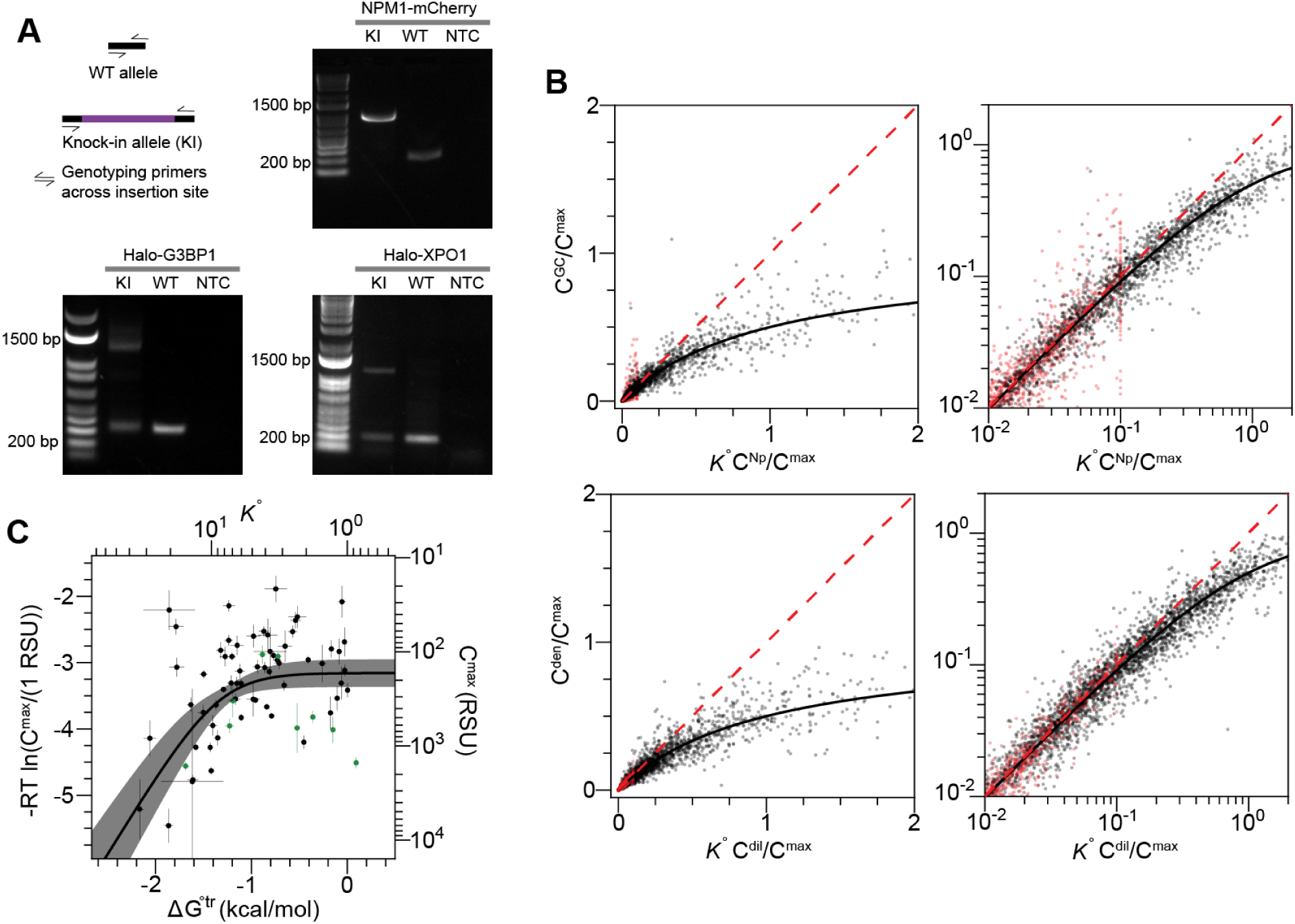
Expression of probes has a hyperbolic dependence. (**A)** Genotyping results for the knock-in cell lines. Schematic of primer design strategy (left); agarose gel electrophoresis with PCR products for the knock-in (KI), wild-type (WT), and no template control (NTC) samples. **(B**) Linear-Linear and Log-Log dimensionless overlay of (top) U2OS GC 148 datafits (5113 data points) and (bottom) other 58 datafits (4424 data points) where x and y axis are normalized by C^max^/K and C^max^, respectively. For those where C^max^ cannot be confidently determined (86 and 20 datasets for A and B, respectively), here distinguished by no measured nucleoplasmic or dilute value greater than 0.1 C^max^/K, those are shown with the max x value normalized to 0.1 and all points red. (**C**) Relationship between partitioning and the max concentration, highlighting that there is a minimal amount of probe accommodated in the GC or stress granule (green points) except at higher partitioning, presumably due to an increase in the number of interactions facilitating phase separation including entry into local environments with smaller mesh sizes.

**Figure S2:**
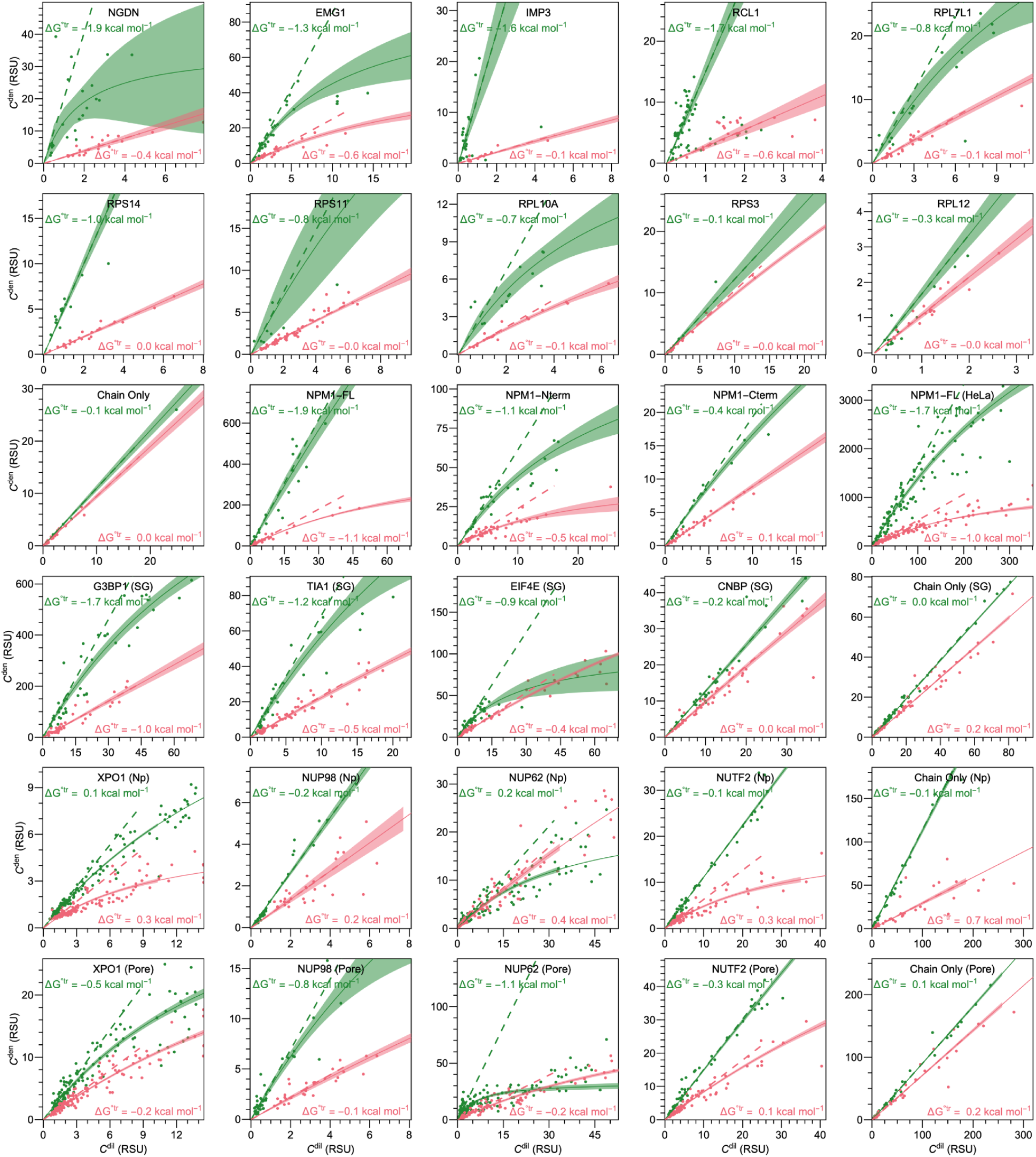
Adding chains to individual constructs always results in LSE. Each construct studied comparing partitioning when fused to no chain (green) and a 400AA chain (red). X axis: C^dil^ (RCU) being Nucleoplasm for GC (no parentheses noted) and Cytoplasm for other compartments listed in parentheses, Y axis: C^den^ (RCU) being the GC (no parentheses noted) or the listed compartments in parentheses.

**Figure S3:**
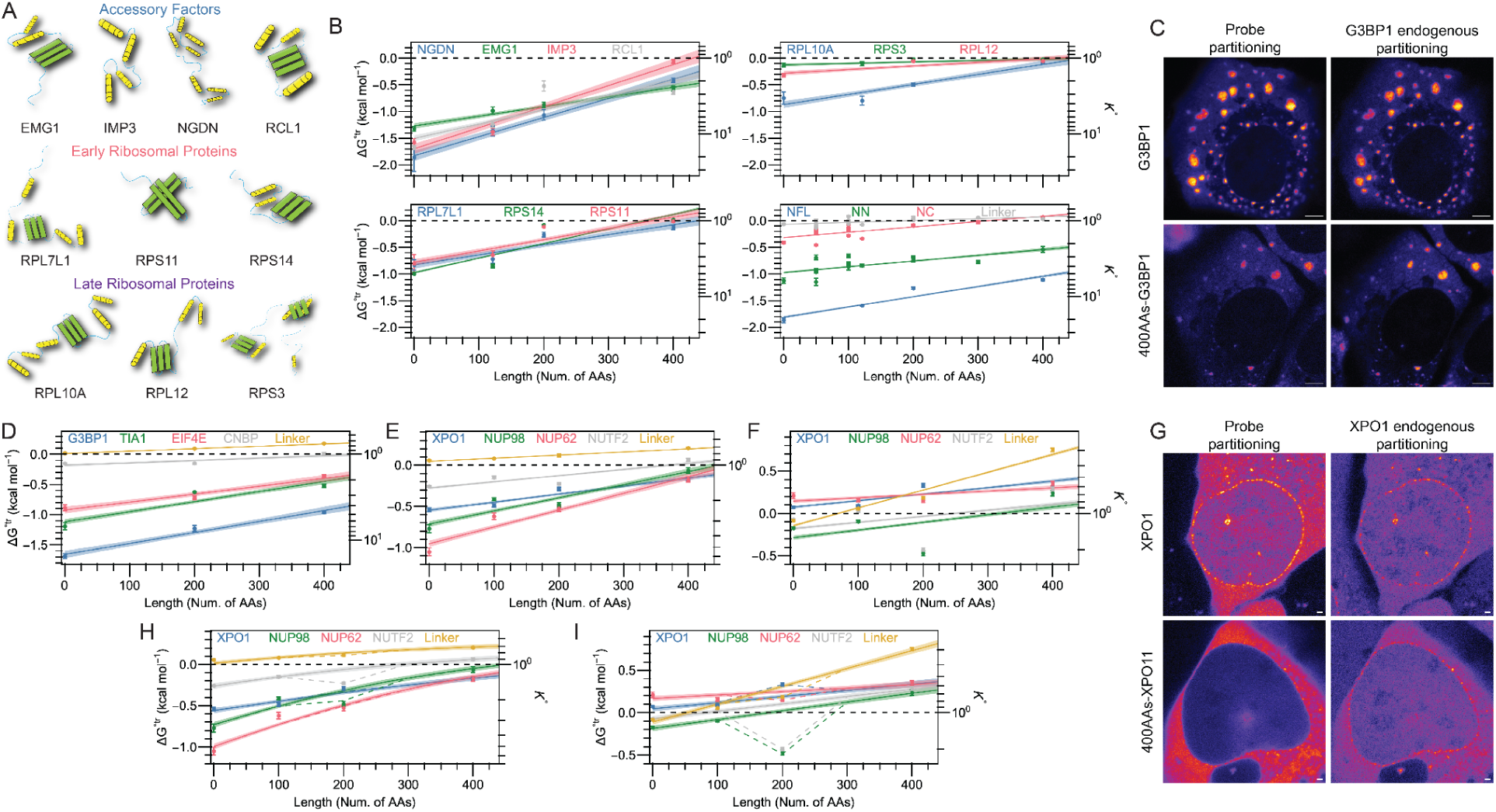
LSE on all constructs. (**A**) Cartoon of nucleolar proteins tested in this manuscript, showing individual structure of each protein. (**B**) Individual Δ*G*^◦^*^tr^*energies with LSE fits for different chain lengths grouped categorically. Plots for GC proteins including accessory factors (top left), early ribosomal proteins (bottom left), late ribosomal proteins (top right), and NPM1 and its truncations (bottom right). Here, ‘linker’ refers to the chain-alone. (**C)** Halo-G3BP1 U2OS cells transfected with G3BP1 probes without and with the 400AAs chain, scale bar 5 µm. **(D)** Stress Granules probe Δ*G*^◦^*^tr^* energies for different chain lengths. **(E)** Nuclear pore proteins LSE fit using a simple linear fit, cytoplasm to nuclear pore. **(F)** Similar to E, but using cytoplasm to nucleoplasm. **(G)** Halo-XPO1 U2OS cells transfected with XPO1 probes, without and with the 400s chain; scale bar: 1 µm. **(H)** Diffraction limit corrected LSE fit, cytoplasm to nuclear pore. **(I)** Diffraction limit corrected LSE fit, cytoplasm to nucleoplasm. In H-I, dashed lines show where the fit adjusts for deviations of the 200-chain length constructs cytoplasmic correction contributes; see SI **12-13**.

**Figure S4:**
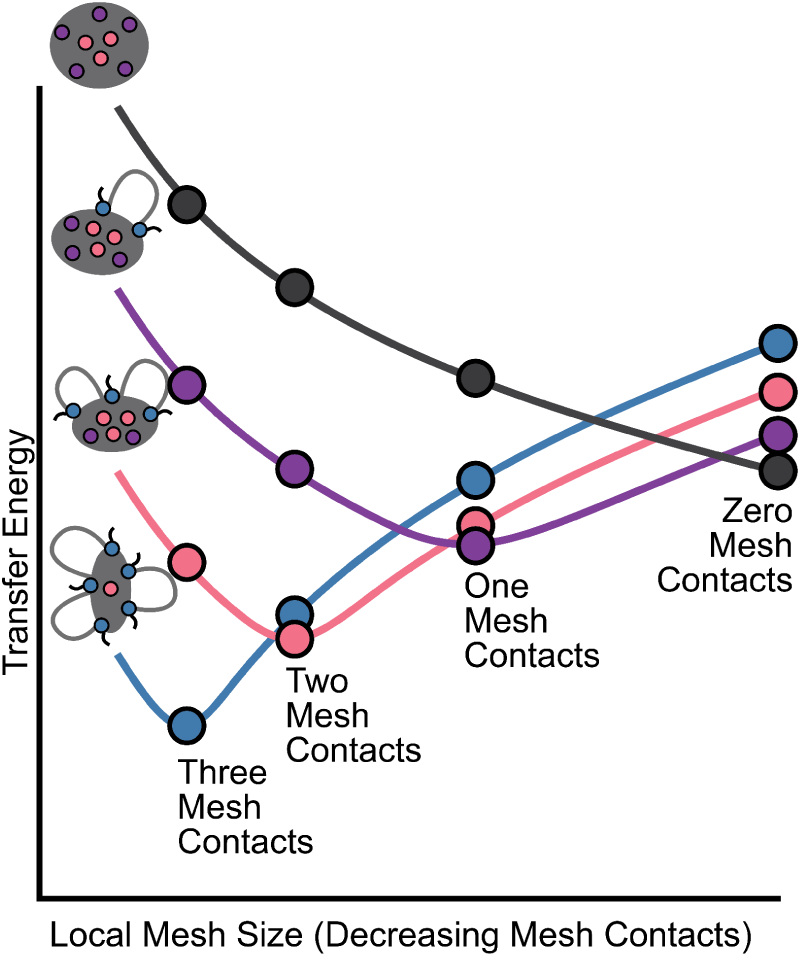
Transfer energy dependence on local mesh size and stage of ribosome biogenesis. The amount of transfer energy needed for contact to be made throughout ribosome biogenesis (blue: accessory factors, pink: early ribosomal proteins, purple: late ribosomal proteins, black: completed ribosome). See methods for full details.

**Figure S5:**
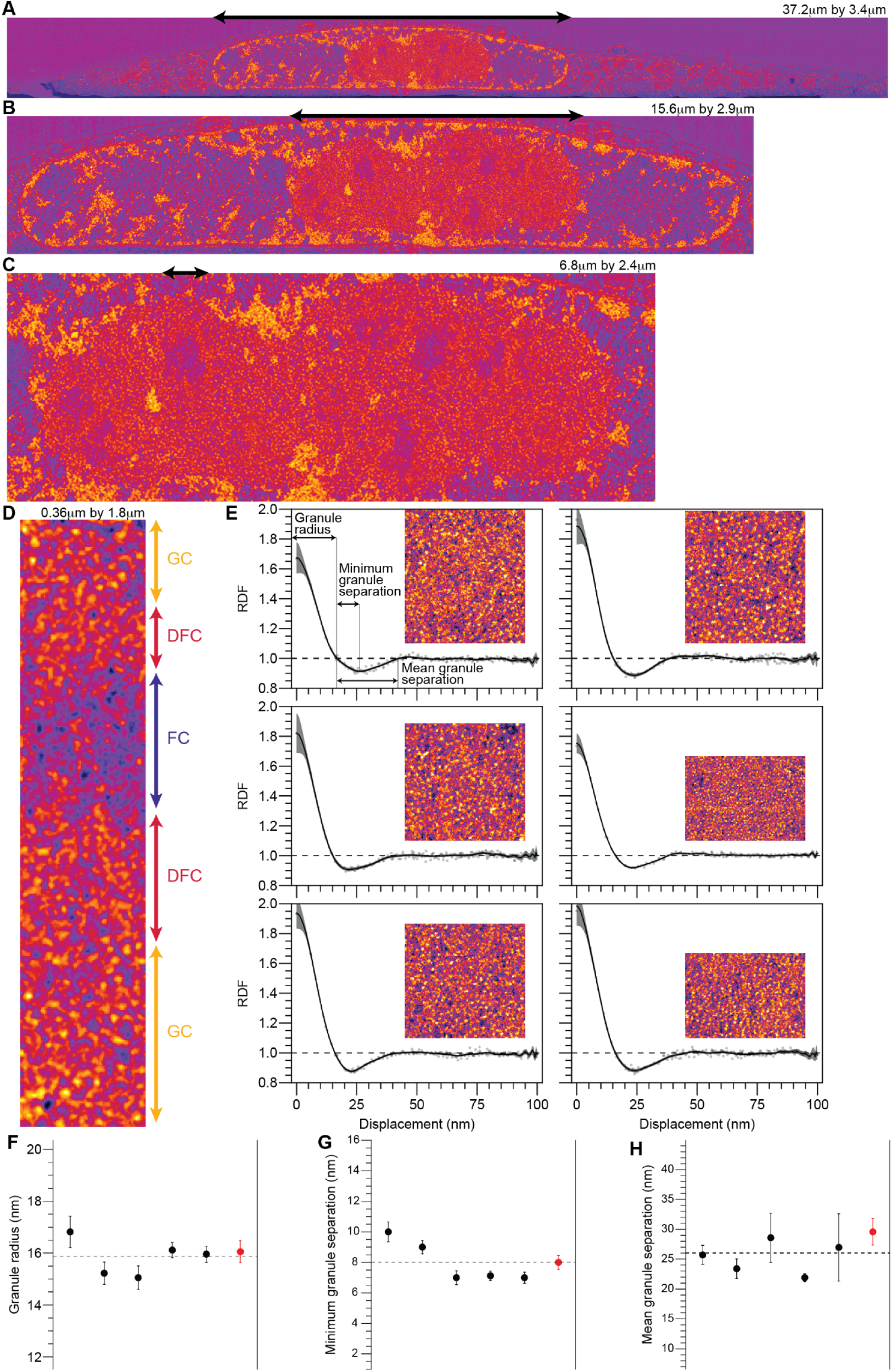
EM data in line with mesh sizes of ribosomal intermediates. (**A-D**) EM data showing subsequently zoomed in regions progressively indicated with arrows and with image sizes as listed. Images slightly blurred for clarity (**D**) Manual delineations of FC, DFC, and GC using a combination of average staining (FC vs. DFC/GC) and the appearance of granules (GC vs. FC/DFC). (**E**) Radial distribution function for GC EM regions (inset) where the first five are from HeLa and the bottom right is Jurkat. Top left plot in **(E)** shows how granule radius, minimum granule separation, and mean granule separation was approximated with each parameter for the six regions shown in (**F**-**H**).

**Figure S6:**
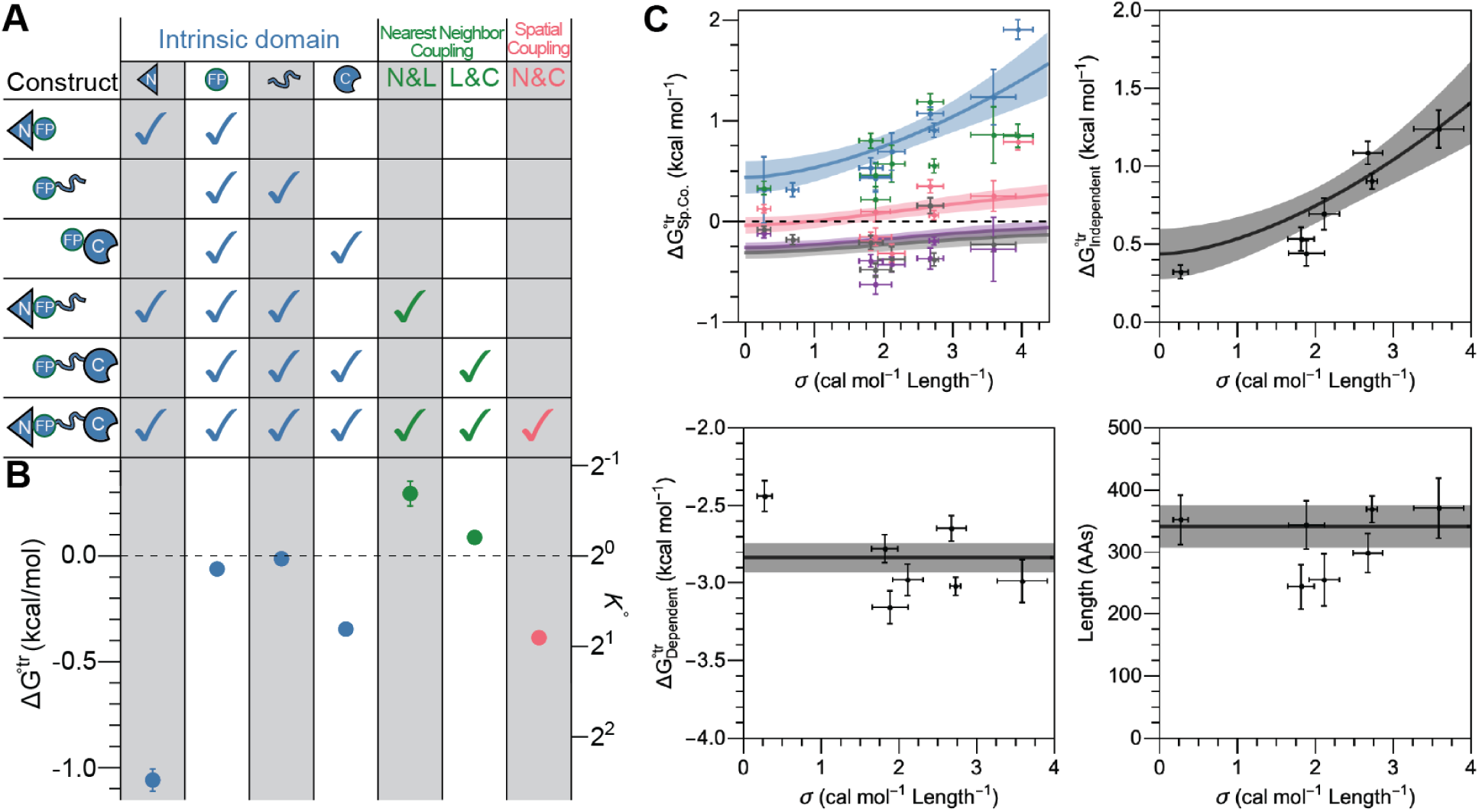
Full LDSC derivation and fit validation. (**A**) Linear breakdown of the Δ*G*^◦^*^tr^* for each NPM1 construct by its domains, including the fluorescent protein’s contributions, into energies as indicated. (**B**) Results of linear decomposition of the Δ*G*^◦^*^tr^*. (**C,** top left) Spatial coupling for NPM1 and ribosome biogenesis factors with chains of length 0 (blue), 121 (red), 200 (gray), and 400 (purple). Additionally, NPM1-N-terminus spatial coupling with 0 chain (green). Individually fit curves shown (top right, and bottom) for specified parameters. Global fit and dependencies are shown in all plots in C.

## Supplementary Tables

**Table S1:**
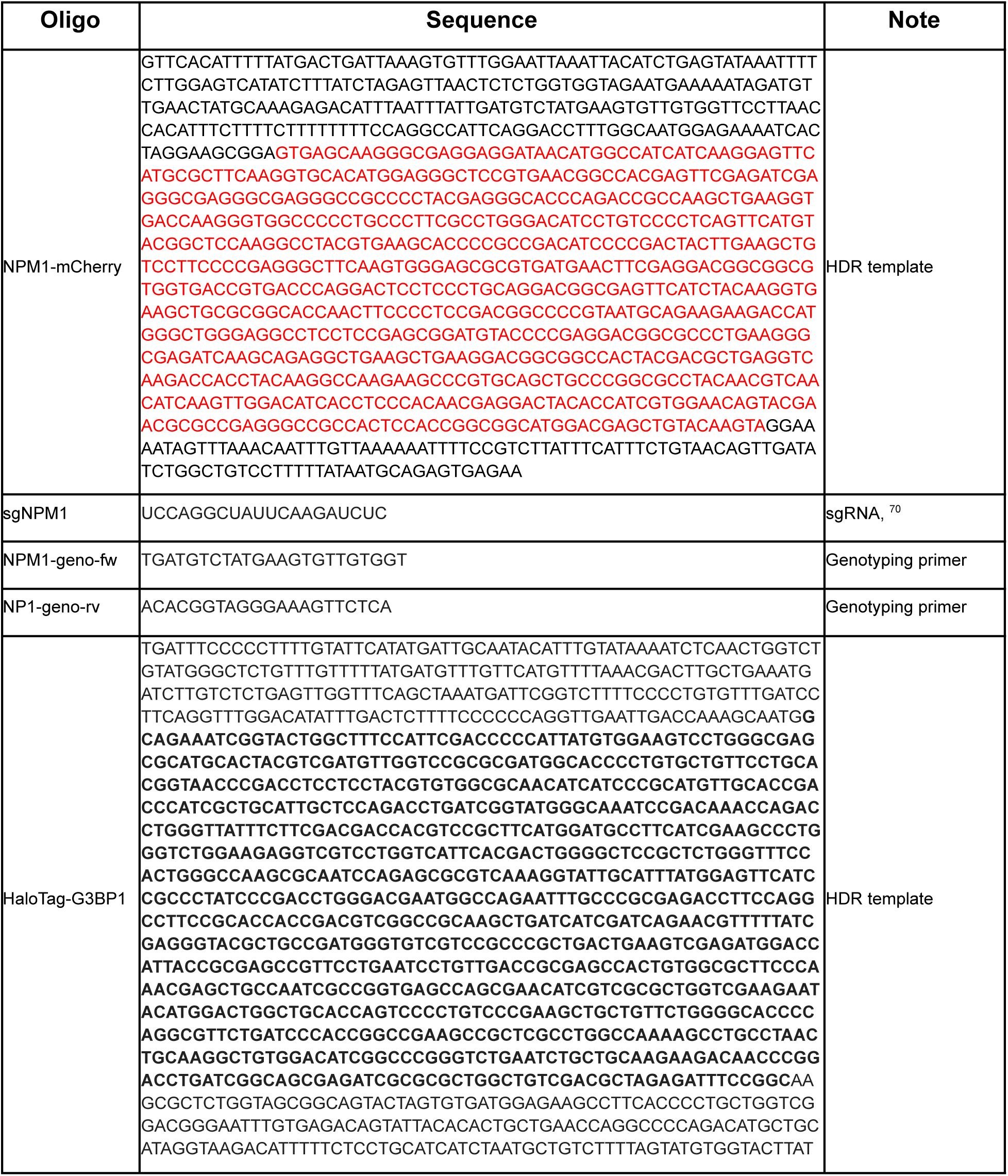
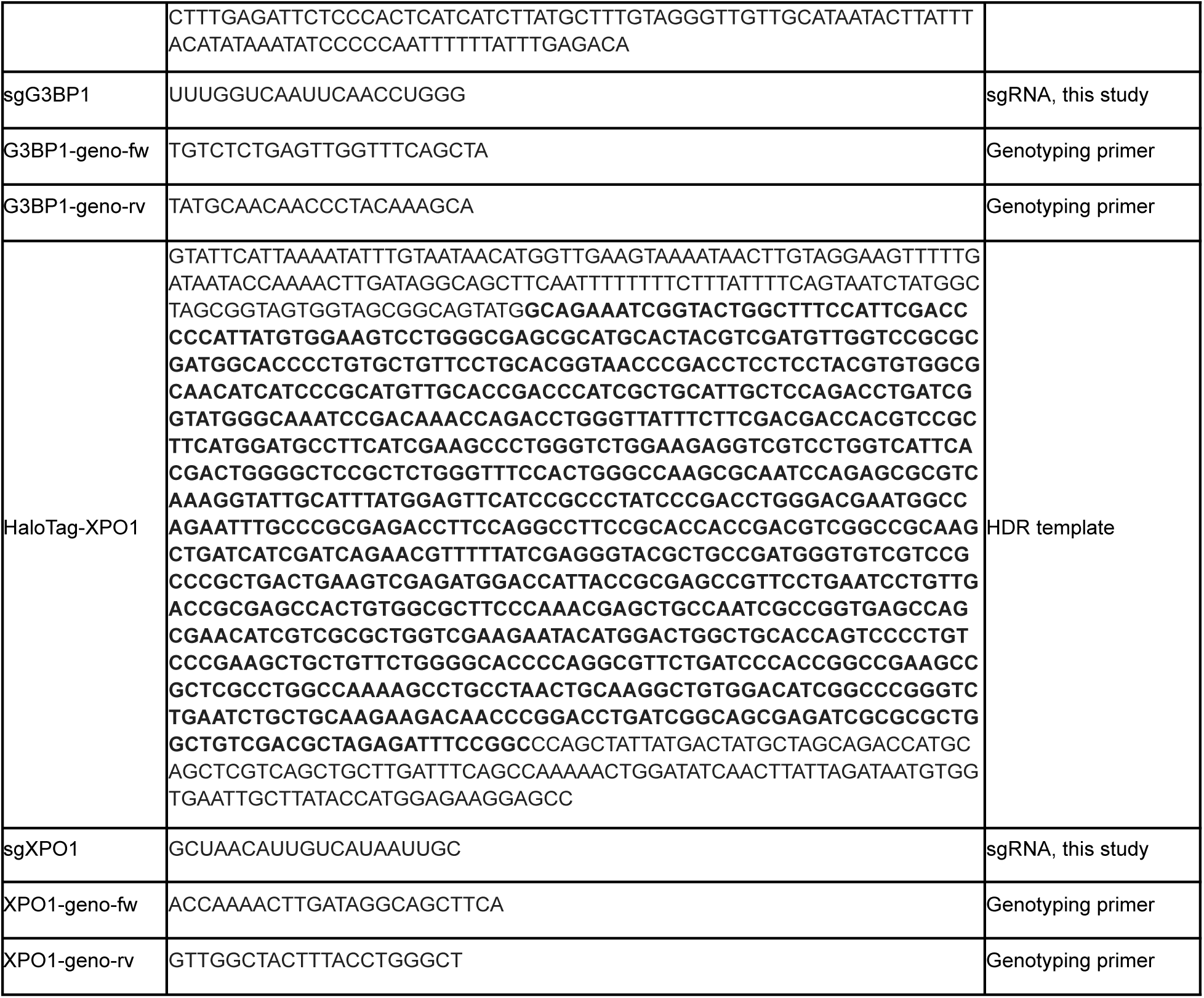
Sequences of HDR template/sgRNA. Nucleotide sequences for CRISPR procedure and validating cell lines after electroporation. Red text is mCherry sequence. Bold is the HaloTag sequence.

**Table S2:**
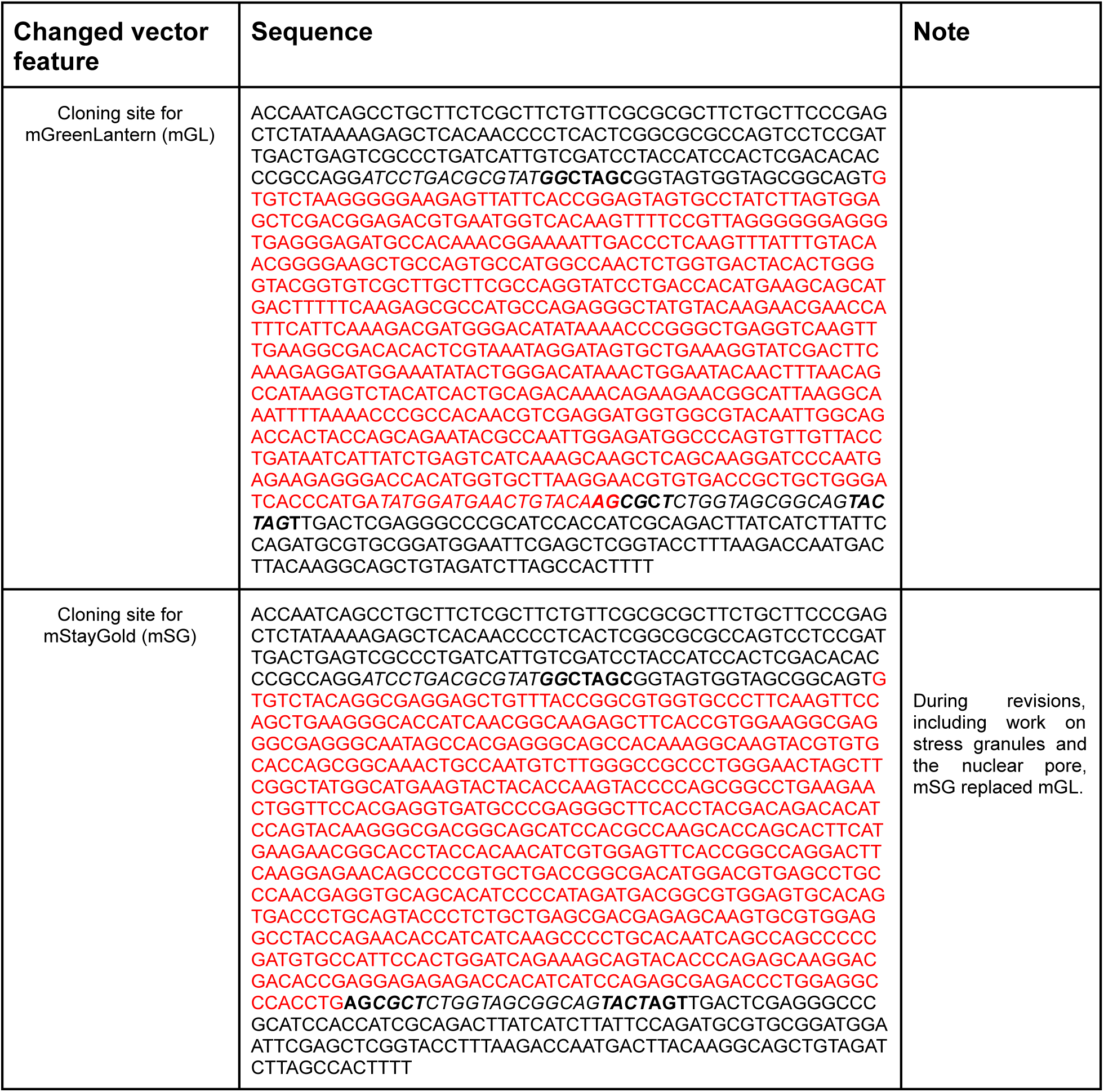
DNA sequence of the vector. Nucleotide sequence and features added to the multicloning vector site. The red bases are mGreenLantern or mStayGold, italicized are the overlap regions and bold are the restriction sites.

**Table S3:**
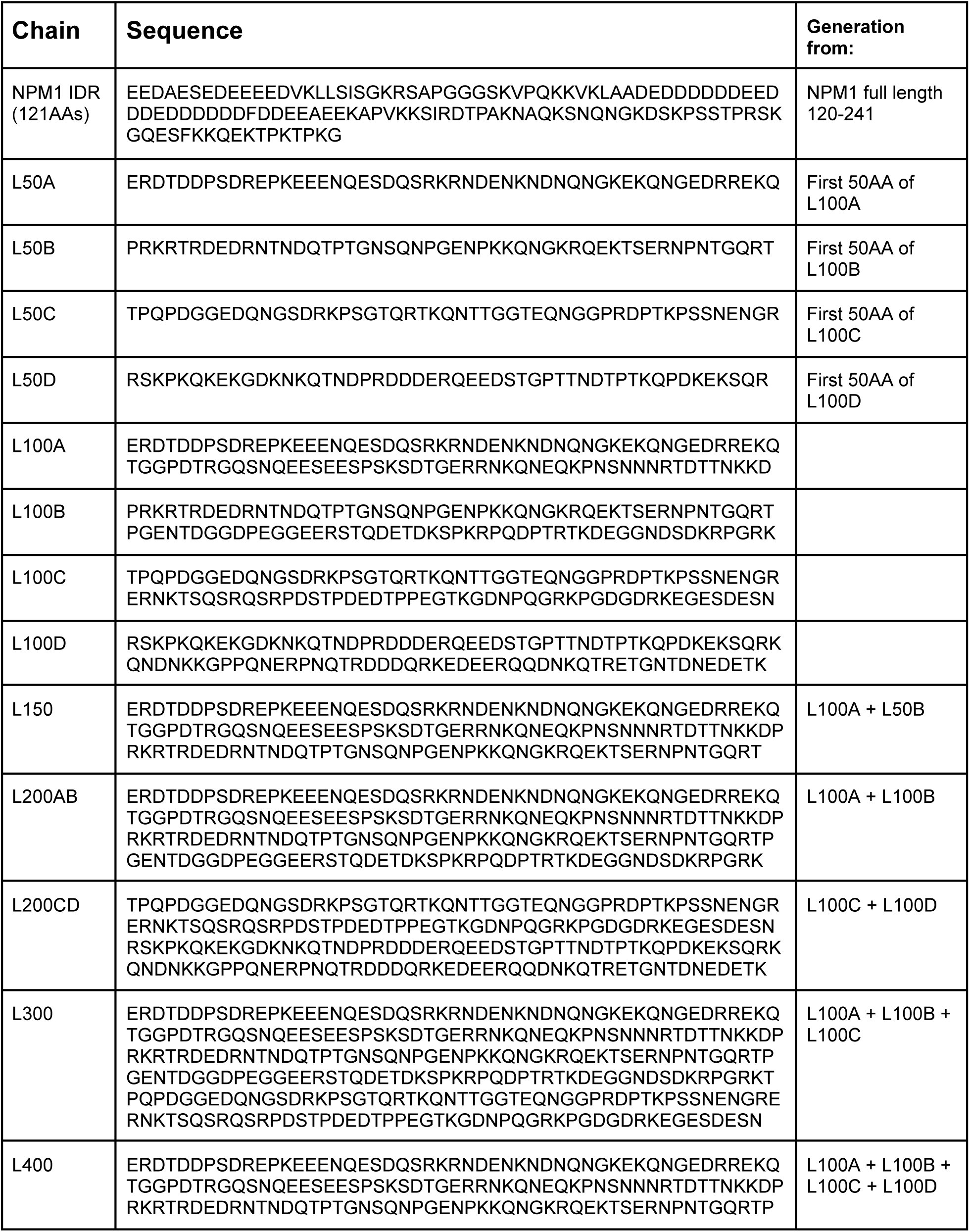
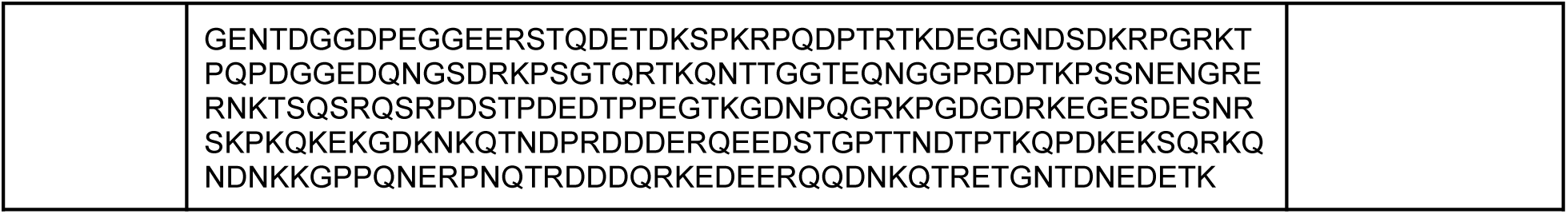
AA Chain sequences. Amino acid sequences for each chain construct. Last column is how longer chains were generated.

**Table S4:**
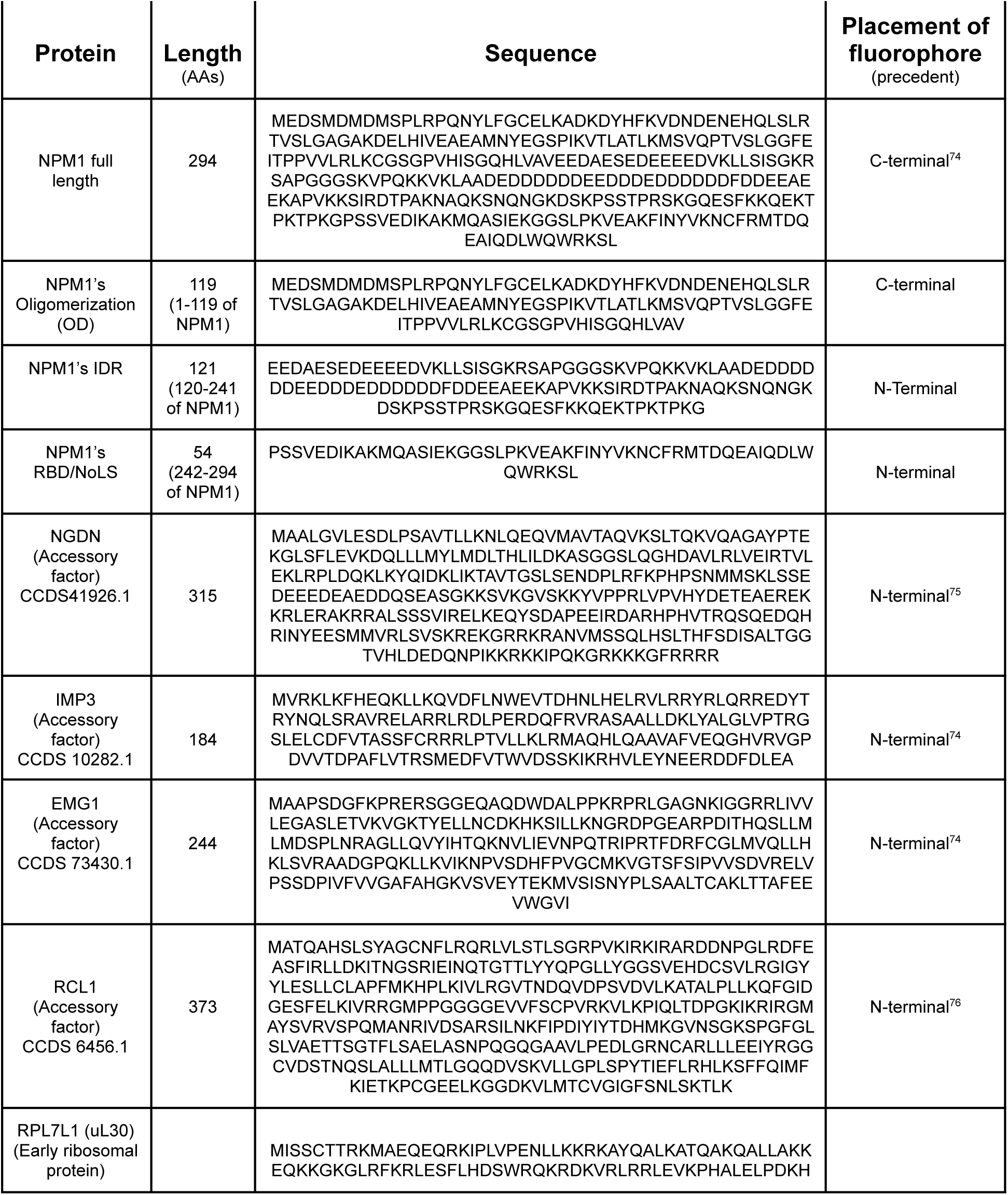
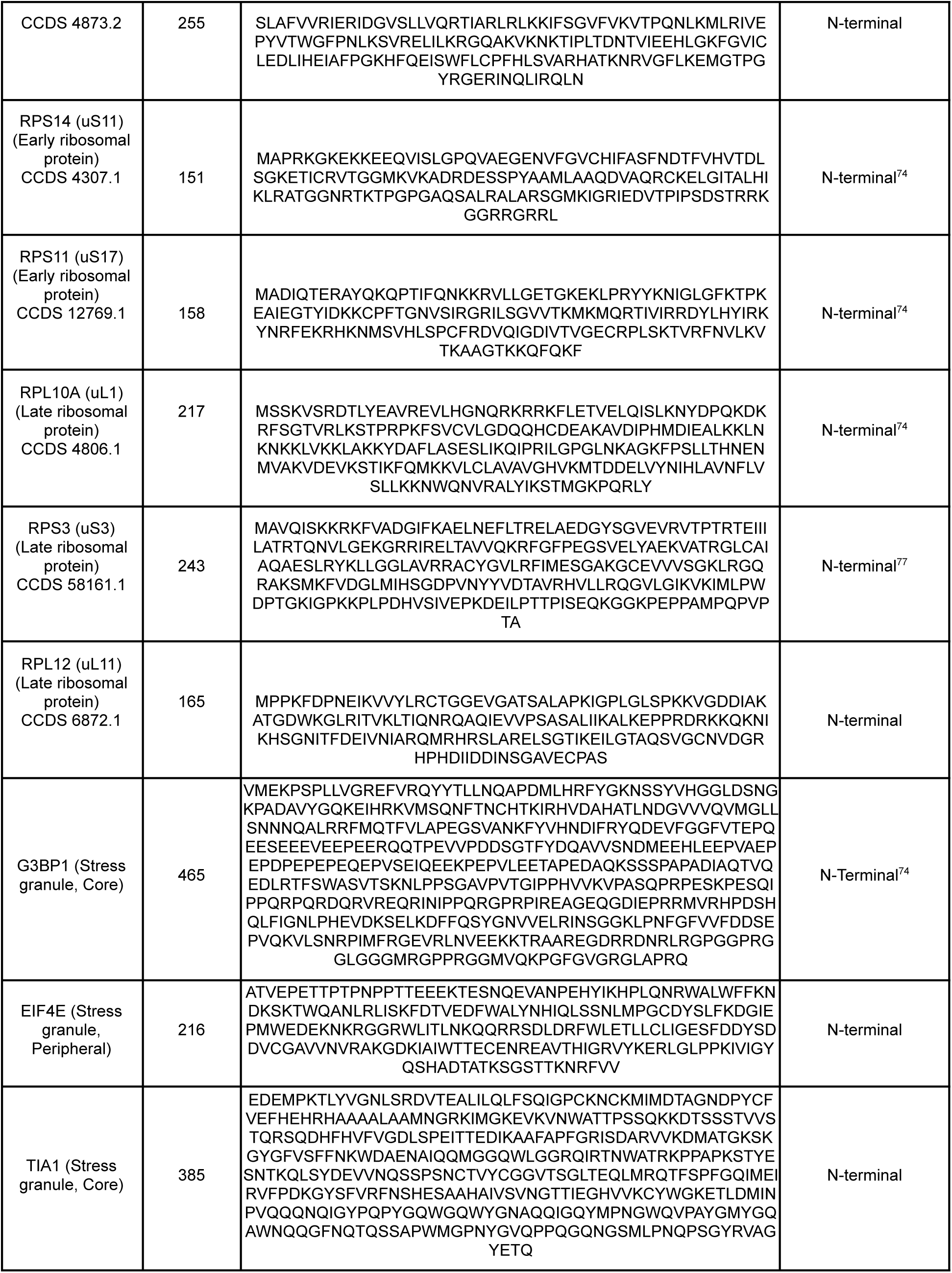
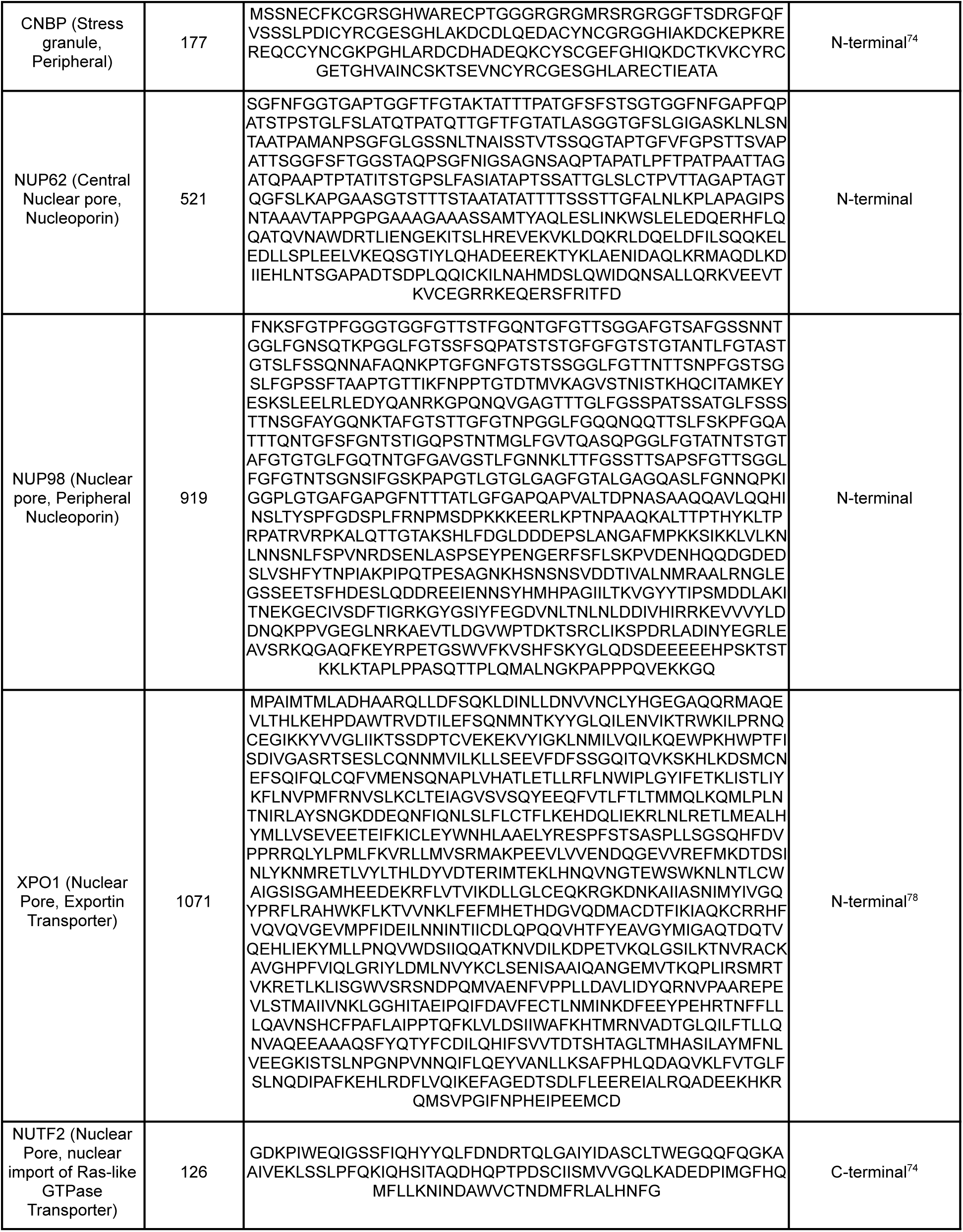
Proteins used in probe design and NPM1 domain truncations. Specific CCDS numbers and ribosome biogenesis roles included for clarity. Proteins for LSE on stress granules and nuclear pore also included. The final column indicates the position of the fluorescent tag relative to the protein of interest, with literature precedent noted where applicable. Some protein inserts do not have an N-terminal methionine, as an upstream one is present (either in the plasmid backbone or the fluorescent tag).

**Table S5.**
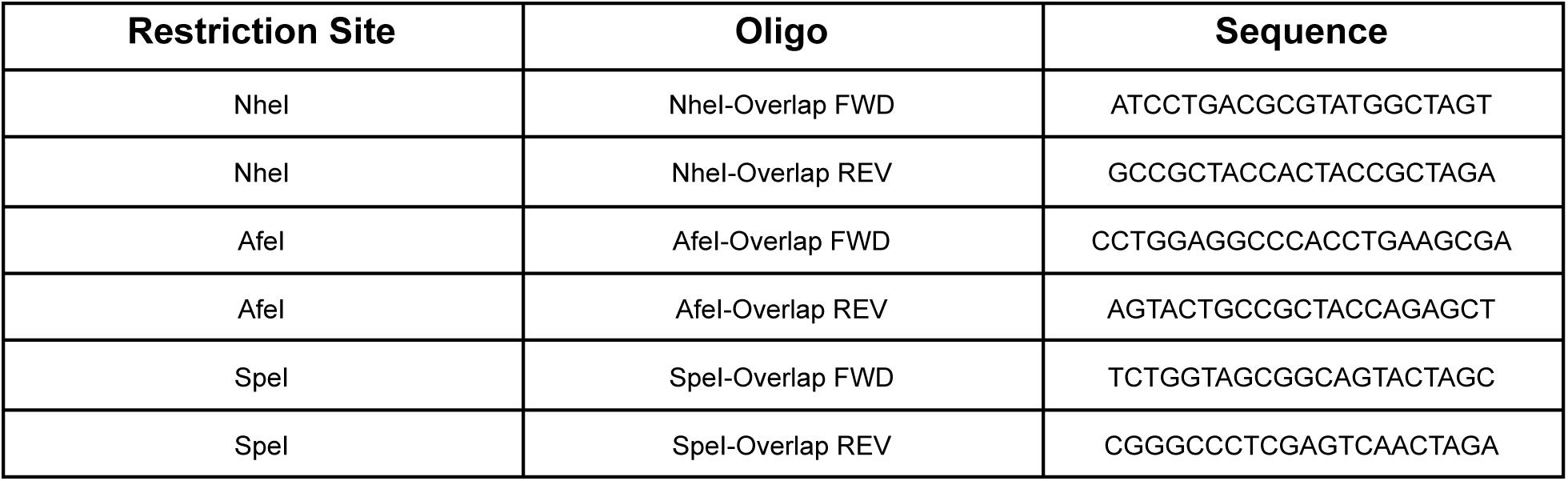
Primer sequences used to generate overlapping homology regions for Gibson Assembly of plasmid constructs. All probes in this study used the same backbone design. Proteins of interest were inserted at either NheI (N-terminal to the fluorophore) or SpeI (C-terminal to the fluorophore), whereas randomized chains were inserted at AfeI (C-terminal to the fluorophore and upstream of SpeI).

**Table S6:** All data points in paper. Cell-level fluorescence measurements for all 186 probes analyzed across three condensates in this study. Each row represents one analyzed cell expressing the indicated probe (plasmid construct). Columns report the condensate, construct abbreviation, probe architecture (N-term, fluorescent protein, chain, and C-term), edited cell line background, and measured signal in the nucleoplasm and/or cytoplasm, and condensate (RCU).

**Table S7:** All. ΔG^°tr^ **and** C^max^ **in paper.** Summary of steady-state transfer free energies (ΔG^°tr^) and C^max^-derived free energies for all probes in this study. Each row corresponds to one probe with columns identifying the condensate, probe name, design (N-term, fluorescent protein, chain, and C-term), which CRISPR-edited cell line was transfected, and the dilute and dense phase regions used for analysis. ΔG^°tr^ reports on the steady-state Transfer Free Energy in kcal/mol with error, whereas “-RT ln(C^max^)” reports the free-energy-transformed C^max^ value in kcal/mol with standard error when available. “Linear?” denotes whether steady-state partitioning was determined from the linear regime (TRUE) or from C^max^ (FALSE) for **Fig. S1**.

**Table S8:** All σ-values (Sigma) in paper. Fitted σ-values and ΔG^°tr^ intercepts for all protein probes and chains analyzed in this study. For each σ-value, fit method (standard fit or Global Fit, **SI 12-13**), fitted σ-value with standard error, and fitted 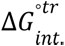 (Intercept Transfer Free Energy) in kcal/mol are listed. For nuclear pore measurements, each is reported for both cytoplasm-to-nucleoplasm and cytoplasm-to-nuclear pore comparisons, with both standard and global fits listed; nuclear pore chains are reported as standard fits only. All nucleolar and stress granule entries use the standard fit.

