## Supplement for "Nanometer condensate organization in live cells derived from partitioning measurements"

*Supplementary Information*

Christina Dollinger[1,4], S. Thomas Hennigan[1,4], Evdokiia Potolitsyna[1,4], Abigail G. Martin[1], Nayara Alcantara-Contessoto[1], Archish Anand[1], Gandhar K. Datar[1,2], Jeremy D. Schmit[3], Joshua A. Riback[1†]

[1]Department of Molecular and Cellular Biology, Baylor College of Medicine; Houston, TX 77030, United States of America
[2]Medical Scientist Training Program, Baylor College of Medicine; Houston, TX 77030, United States of America
[3]Department of Physics, Kansas State University, Manhattan, KS 66506, United States of America
[4]These authors contributed equally.

### Contents

**1 A note of caution on the use of the term standard state transfer free energy. 3**

**2 Extraction of the steady-state transfer free energy 3**

**3 Expression dependent partitioning 4**

**4 Basic statistical mechanics of the transfer free energy 4**

**5 Theoretical basis for the $\sigma$-value approach 6**

**6 Local Size Exclusion 7**

**7 Elucidating microenvironment heterogeneity via $\sigma$-value and LSE 11**

**8 Model for local energetics as a function of mesh size during ribosome biogenesis 11**

**9 Domain decomposition and nearest-neighbor coupling 13**

**10 Theoretical basis for Length-Dependent Spatial Coupling 15**

**11 Implementation of the linear decomposition 17**

**12 Theoretical basis for studying small condensates 18**

**13 Diffraction-limited model for nuclear pore measurements 19**

#### 1 A note of caution on the use of the term standard state transfer free energy.

We use the superscript $\circ$ in $K^{\circ}$ and $\Delta G^{\circ \mathrm{tr}}$ to denote an *apparent standard state* defined by the measured steady-state partition coefficient and the corresponding transfer free energy, respectively, of the constructs between the dilute and dense cellular environments (e.g., nucleoplasm and nucleolus) under unperturbed interphase conditions. Thus, this notation refers to measurements obtained under baseline cellular conditions, without induced overexpression, and averaged across the interphase cell population at steady state. The application of standard-state terminology from physical chemistry should be considered with some caution, especially in light of previous debates on the use of such terms[1, 2]. Indeed, the notion that the regions studied are true phases and perfectly in a steady state is likely incorrect to some extent[3, 4]. Furthermore, interpretation of our measurements largely assumes a single dense phase within a marker-defined compartment; if future work demonstrates that multiple unresolved dense phases contribute substantially to the measured signal, this would complicate the interpretation of the inferred thermodynamic quantities. Although additional corrections may ultimately refine these quantities to more rigorous thermodynamic descriptions, we find that this framework provides a useful, compact way to interpret partitioning measurements of constructs.

#### 2 Extraction of the steady-state transfer free energy

To determine the steady-state $\Delta G^{\circ \mathrm{tr}}$ for a specific probe, single-cell measurements of the dense phase ($C_{\mathrm{den}}$) and dilute phase ($C_{\mathrm{dil}}$) concentrations, were empirically fit to:

$$C_{\mathrm{den}} = \frac{C_{\mathrm{dil}}}{a + bC_{\mathrm{dil}}} \tag{S1}$$

noting that this equation corresponds to a rectangular hyperbola as it can be arranged into the form:

$$\left(C_{\mathrm{dil}} + \frac{a}{b}\right)\left(C_{\mathrm{den}} - \frac{1}{b}\right) = -\frac{a}{b^2} \tag{S2}$$

and where the parameters are defined as:

$$a = \frac{1}{K^{\circ \mathrm{tr}}} \tag{S3A}$$

$$= e^{\frac{\Delta G^{\circ \mathrm{tr}}}{RT}} \tag{S3B}$$

and

$$b = \frac{1}{C_{\mathrm{den}}^{\mathrm{max}}} \tag{S4A}$$

$$= e^{\frac{\Delta G^{c\,\mathrm{max}}}{RT}} \tag{S4B}$$

Where the $\Delta G^{c\,\mathrm{max}}$ is referenced to RCU units and intended primarily to enforce the constraint $b > 0$.

This form (equation S1) was chosen because it captures the experimentally observed limits of low-expression partitioning and high-expression saturation:

$$\lim_{C_{\mathrm{dil}} \to 0} C_{\mathrm{den}} = K^{\circ \mathrm{tr}}\, C_{\mathrm{dil}} \tag{S5A}$$

$$\lim_{C_{\mathrm{dil}} \to \infty} C_{\mathrm{den}} = C_{\mathrm{den}}^{\mathrm{max}} \tag{S5B}$$

### 3 Expression dependent partitioning

All constructs show clear hyperbolic expression-dependent partitioning. To rationalize this, we invoke the concept that there is a maximum concentration of available sites within the condensate to which a probe can localize to, $C_{\text{den}}^{\text{max}}$ with free and occupied sites being $S_{\text{den}}^{\text{free}}$ and $S_{\text{den}}^{\text{occ}}$, respectively. Additionally, we define $k_{a,\text{dil}}$ as the occupancy equilibrium per concentration of sites. Thus, the localization sites within the condensate can be written:

$$C_{\text{den}}^{\text{max}} = S_{\text{den}}^{\text{free}} + S_{\text{den}}^{\text{occ}} \tag{S6A}$$

$$= S_{\text{den}}^{\text{free}} + k_{a,\text{dil}} C_{\text{dil}} S_{\text{den}}^{\text{free}} \tag{S6B}$$

$$= S_{\text{den}}^{\text{free}} \left(1 + k_{a,\text{dil}} C_{\text{dil}}\right) \tag{S6C}$$

Rearranging equation S6C for $S_{\text{den}}^{\text{free}}$ yields:

$$S_{\text{den}}^{\text{free}} = \frac{C_{\text{den}}^{\text{max}}}{1 + k_{a,\text{dil}} C_{\text{dil}}} \tag{S7}$$

Employing S7, we can now solve for $C_{\text{den}}$:

$$C_{\text{den}} = S_{\text{den}}^{\text{occ}} \tag{S8A}$$

$$= k_{a,\text{dil}} C_{\text{dil}} S_{\text{den}}^{\text{free}} \tag{S8B}$$

$$= k_{a,\text{dil}} C_{\text{dil}} \frac{C_{\text{den}}^{\text{max}}}{1 + k_{a,\text{dil}} C_{\text{dil}}} \tag{S8C}$$

$$= \frac{C_{\text{dil}}}{\frac{1}{k_{a,\text{dil}} C_{\text{den}}^{\text{max}}} + \frac{C_{\text{dil}}}{C_{\text{den}}^{\text{max}}}} \tag{S8D}$$

$$= \frac{C_{\text{dil}}}{\frac{1}{K^{\text{otr}}} + \frac{C_{\text{dil}}}{C_{\text{den}}^{\text{max}}}} \tag{S8E}$$

where, by equation S5A, $K^{\text{otr}} = k_{a,\text{dil}} C_{\text{den}}^{\text{max}}$.

This description should be viewed as a coarse-grained model; the “sites” need not correspond to discrete binding pockets but instead represent the finite number of favorable environments available within the condensate per average volume of the dense phase.

### 4 Basic statistical mechanics of the transfer free energy

To interpret $\Delta G^{\text{otr}}$ more generally, we consider the Widom insertion of a probe construct into a background ensemble of microstates describing each phase[5]. Let $\Omega$ denote a microstate of phase $i$. The energy associated with this configuration is $g_0^i(\Omega)$ and its corresponding Boltzmann weight is

$$w^i(\Omega) = e^{-\frac{g_0^i(\Omega)}{RT}}. \tag{S9}$$

The probability of observing configuration $\Omega$ is therefore

$$P^i(\Omega) = \frac{e^{-\frac{g_0^i(\Omega)}{RT}}}{Z_0^i} \tag{S10A}$$

$$= \frac{w^i(\Omega)}{Z_0^i}, \tag{S10B}$$

where the partition function of the phase is

$$Z_0^i = \int w^i(\Omega)\, d\Omega \tag{S11}$$

and the total Gibbs free energy of the phase is

$$G_0^i = -RT \ln Z_0^i. \tag{S12}$$

To compute the free energy associated with inserting a probe construct $\alpha$, we consider placing the probe with configuration $\Gamma$ within a background microstate $\Omega$. The Boltzmann weight associated with this insertion is

$$w^i(\alpha, \Gamma|\Omega) = e^{-\frac{g^i_\alpha(\Omega,\Gamma)}{RT}}. \tag{S13}$$

Integrating over the probe's internal degrees of freedom yields the effective insertion weight, which represents the Boltzmann weight for inserting the probe into the background microstate $\Omega$.

$$w^i(\alpha|\Omega) = \int w^i(\alpha, \Gamma|\Omega)\, d\Gamma \tag{S14A}$$

$$\equiv e^{-\frac{g^i_\alpha(\Omega)}{RT}}. \tag{S14B}$$

The partition function of the system containing the probe is therefore

$$Z_\alpha^i = \int w^i(\Omega) w^i(\alpha|\Omega)\, d\Omega \tag{S15A}$$

$$= \int Z_0^i P^i(\Omega) w^i(\alpha|\Omega)\, d\Omega \tag{S15B}$$

$$= Z_0^i \int P^i(\Omega) w^i(\alpha|\Omega)\, d\Omega \tag{S15C}$$

$$= Z_0^i \left\langle w(\alpha) \right\rangle_0^i. \tag{S15D}$$

Here, we have defined the average insertion weight

$$\left\langle w(\alpha) \right\rangle_0^i = \int P^i(\Omega) w^i(\alpha|\Omega)\, d\Omega, \tag{S16}$$

which represents the Boltzmann weight of inserting $\alpha$ averaged over the configurations of the phase. The corresponding insertion free energy, equivalent to the excess chemical potential of the probe at infinite dilution,

$$G_\alpha^{\mathrm{ex},i} = G_\alpha^i - G_0^i \tag{S17A}$$

$$= -RT \ln \frac{Z_\alpha^i}{Z_0^i} \tag{S17B}$$

$$= -RT \ln \left\langle w(\alpha) \right\rangle_0^i. \tag{S17C}$$

The transfer free energy between dilute and dense phases for probe $\alpha$ can therefore be written

$$\Delta G_\alpha^{\mathrm{otr}} = G_\alpha^{\mathrm{ex,den}} - G_\alpha^{\mathrm{ex,dil}} \tag{S18A}$$

$$= -RT \ln \frac{\left\langle w(\alpha) \right\rangle_0^{\mathrm{den}}}{\left\langle w(\alpha) \right\rangle_0^{\mathrm{dil}}} \tag{S18B}$$

$$\approx -RT \ln \left\langle w(\alpha) \right\rangle_0^{\mathrm{den}}. \tag{S18C}$$

The final approximation assumes that interactions experienced by the probe in the dilute phase are negligible relative to those in the dense phase, such that $\langle w(\alpha)\rangle_0^{\text{dil}} \approx 1$. In this limit, the transfer free energy is dominated by interactions experienced by the probe within the dense phase. More generally, the equation S18C gives us a theoretical framework to understand how partitioning measurements are determined by the ensemble of local microstates within a condensate.

### 5 Theoretical basis for the $\sigma$-value approach

We now extend the framework above to consider a probe construct $\alpha - \beta$, where $\beta$ represents a chain of $N$ residues appended to $\alpha$. The partition function of this composite probe can be written

$$Z^i_{\alpha-\beta} = \int w^i(\Omega) w^i(\alpha|\Omega) w^i(\beta|\Omega,\alpha)\, d\Omega. \tag{S19}$$

Using Bayes' theorem and Eq. S15D, the conditional probability of configuration $\Omega$ given that $\alpha$ is present is

$$P^i(\Omega|\alpha) = P^i(\Omega)\frac{w^i(\alpha|\Omega)}{\langle w(\alpha)\rangle_0^i} \tag{S20A}$$

$$= \frac{w^i(\Omega)}{Z^i_0}\frac{w^i(\alpha|\Omega)}{\langle w(\alpha)\rangle_0^i} \tag{S20B}$$

$$= \frac{w^i(\Omega) w^i(\alpha|\Omega)}{Z^i_\alpha}. \tag{S20C}$$

Substituting this expression into Eq. S19 yields

$$Z^i_{\alpha-\beta} = Z^i_\alpha \int P^i(\Omega|\alpha) w^i(\beta|\Omega,\alpha)\, d\Omega \tag{S21A}$$

$$= Z^i_\alpha \langle w(\beta)\rangle^i_\alpha, \tag{S21B}$$

where the average insertion weight of $\beta$ conditioned on $\alpha$ is

$$\langle w(\beta)\rangle^i_\alpha = \int P^i(\Omega|\alpha) w^i(\beta|\Omega,\alpha)\, d\Omega. \tag{S22}$$

The free energy of the composite probe therefore becomes

$$G^i_{\alpha-\beta} = -RT\ln Z^i_{\alpha-\beta} \tag{S23A}$$

$$= -RT\ln Z^i_\alpha - RT\ln\langle w(\beta)\rangle^i_\alpha \tag{S23B}$$

$$= G^i_\alpha - RT\ln\langle w(\beta)\rangle^i_\alpha. \tag{S23C}$$

The quantity $-RT\ln\langle w(\beta)\rangle^i_\alpha$ represents the free energy cost of appending a chain $\beta$ to a probe $\alpha$ that is already present in phase $i$. Because $\beta$ consists of a chain of $N$ residues, we write

$$\langle w(\beta)\rangle^i_\alpha = \langle w(N)\rangle^i_\alpha. \tag{S24}$$

Over the experimentally sampled range of chain lengths, the effect of increasing the chain length is linear. Therefore, the insertion free energy can be expanded to the first order in chain length $N$:

$$-RT \ln \langle w(N) \rangle_\alpha^i = -RT \ln \langle w(N) \rangle_\alpha^i \Big|_{N=0} - NRT \left. \frac{\langle w'(N) \rangle_\alpha^i}{\langle w(N) \rangle_\alpha^i} \right|_{N=0} + \dots \tag{S25A}$$

$$= -NRT \langle w'(0) \rangle_\alpha^i + \dots \tag{S25B}$$

Evaluating the derivative yields

$$-NRT \langle w'(0) \rangle_\alpha^i = -NRT \int P^i(\Omega|\alpha) \left. \frac{d}{dN} e^{-\frac{g(N|\Omega,\alpha)}{RT}} \right|_{N=0} d\Omega \tag{S26A}$$

$$= N \int P^i(\Omega|\alpha) \left. \frac{dg(N|\Omega,\alpha)}{dN} \right|_{N=0} d\Omega. \tag{S26B}$$

where the second line uses the fact that $g(0|\Omega, \alpha) = 0$. Defining

$$\sigma(\Omega, \alpha) = \left. \frac{dg(N|\Omega,\alpha)}{dN} \right|_{N=0}, \tag{S27}$$

and averaging over configurations gives

$$\sigma_\alpha^i = \int P^i(\Omega|\alpha) \sigma(\Omega, \alpha) \, d\Omega. \tag{S28}$$

Substituting into Eq. S23C yields the approximate result

$$G_{\alpha-\beta}^i \approx G_\alpha^i + N\sigma_\alpha^i. \tag{S29}$$

Thus, $\sigma_\alpha^i$ represents the marginal free energy cost of extending the chain by one residue within phase $i$. Analogous to other derivatives of the Gibbs free energy (e.g., the chemical potential), $\sigma$ reflects the average interaction free energy experienced by an additional residue within the environment of the phase. In general, this quantity may contain contributions from multiple physical effects, including confinement, electrostatic interactions, and specific chemical interactions. In section 6 we introduce a simplified description in which $\sigma$ is dominated by confinement imposed by the local mesh size of the condensate, which we term Local Size Exclusion (LSE). In section 7, we make some points about the utility of the $\sigma$-value in the context of inferring heterogeneity of condensate microenvironments.

### 6 Local Size Exclusion

For the purposes of this manuscript, we approximate the general $\sigma$-value as being dominated by confinement of the appended chain within the meshwork of the condensate locally around protein $\alpha$. We will refer to this measurement as Local Size Exclusion (LSE), as the mathematics is analogous to that employed in understanding size exclusion chromatography [6, 7, 8]. To do this, we utilize a random, non-hydrophobic chain (no aromatic or aliphatic residues, e.g. only R,K,D,E,Q,N,S,T,P, and G) leading to the approximation:

$$\sigma_\alpha^i \approx \sigma_{\alpha,\mathrm{LSE}}^i \tag{S30}$$

Because the appended chains are sequence-randomized, they are fairly unique compared to the grammar of disordered regions in proteins. As an additional check, we analyzed the randomized chains with

NARDINI+[9] to test for unintended IDR-like patterning. The chains showed little evidence of strong patterning features, consistent with their randomized design. In our experiments, the dependence of the transfer free energy on chain length when fused to a $\alpha$ is well described by the linear relation:

$$\Delta G^{\text{otr}}(N) = \Delta G^{\text{otr}}_{\text{int.}} + N\sigma_{\text{LSE}} \tag{S31}$$

where $\Delta G^{\text{otr}}_{\text{int.}}$ is the linear extrapolation to zero chain (the subscript denotes "intercept") and $\sigma_{\text{LSE}}$ is the LSE $\sigma$-value. We note that to simplify notation and in conceptual spirit with equation S18C, we have used $\sigma$ rather than the more technically correct $\Delta\sigma = \sigma^{den} - \sigma^{dil}$. In future work, where neither phase can be disregarded with respect to the chain, then it may be more appropriate to write $\Delta\sigma$.

To convert the LSE $\sigma$-value into an effective local mesh size, we use the scaling relation for the confinement free energy of a polymer in a region of characteristic dimension $D$. The length $N$ chain is split into $N/g$ compression blobs, where $g$ is the number of monomers per compression blob. The blob size is determined by the number of monomers where the confinement free energy is approximately equal to the thermal energy randomizing the configuration of the chain, or $\Delta G^{\text{tr}}_{\text{confinement}} \approx RTN/g$ [10, 7]. For confinement in one or two dimensions, this yields the relationship $D = R_{\text{ee},0}g^{\nu}$ where $R_{\text{ee},0}$ is the end-to-end distance prefactor and $\nu$ is the flory exponent that depends on solvent quality[10]. This yields the following conversions:

$$N\sigma_{\text{LSE}} = \Delta G^{\text{tr}}_{\text{confinement}} \tag{S32A}$$

$$\approx RT\frac{N}{g} \tag{S32B}$$

$$= RT\frac{N}{(D/R_{\text{ee},0})^{1/\nu}} \tag{S32C}$$

Simple rearrangement yields the employed conversion:

$$D \approx R_{\text{ee},0}\left(\frac{RT}{\sigma_{\text{LSE}}}\right)^{\nu} \tag{S33}$$

Where it is noted that (1) the proportionality $\Delta G^{\text{tr}}_{\text{confinement}} \approx RT(N/g)$ has a near unity prefactor dependent on geometrical features of the confinement and (2) only is linear with N in the case of narrow confinement $g \ll N$ [10]. For these two reasons, we refer to the evaluation of this equation as the 'effective' local mesh size. Additional error comes from (3) our use of scaling laws for unfolded proteins in water ($R_{ee,0} = 0.55$ nm, $\nu = 0.55$)[11]. To assess the impact of these three points on our ability to quantitatively measure mesh size, we discuss them at length below:

#### 6.1 Prefactor

Here, we use a unity prefactor in the relationship $\Delta G^{\text{tr}}_{\text{confinement}} \propto RT(N/g)$. Simple and calculable geometric models for confinement of a random walk in between a slab, cylinder, and sphere yield a prefactor that is approximately 1.6, 3.9, and 6.6, respectively (where here N/g is referenced to the end-to-end distance of the random walk and the diameter of confinement, being exactly $\frac{1}{6}\pi^2$, $\frac{4}{6}\beta_1^2$, and $\frac{4}{6}\pi^2$, [8]). Note that as the confinement dimensions increase, the prefactor increases. Typically, the prefactor of confinement in a slit, often referred to as the Casassa equation, is employed[8, 7]. However, weak gels are unlikely to be as rigid, with recent results suggesting a universal equation for partitioning into flexible polymer networks $\Delta G^{\text{tr}}_{\text{confinement}} = 4RT\left(\frac{R_g}{l_{\text{cycle}}}\right)^2$ that is applicable to various polymer network densities and topologies[12]. The length scale $l_{\text{cycle}}$ is the mean distance between centroids of elastically effective cycles. To convert

$l_{\text{cycle}}$ into a mesh diameter ($D$), we equate the effective volume of the elastic cycle into an apparent average sphere such that:

$$l_{\text{cycle}} = V_{\text{cycle}}^{1/3} \tag{S34A}$$

$$= \left(\frac{\pi}{6}D^3\right)^{1/3} \tag{S34B}$$

$$= \left(\frac{\pi}{6}\right)^{1/3} D \tag{S34C}$$

Thus, the confinement free energy becomes:

$$\Delta G^{\text{tr}}_{\text{confinement}} = 4RT\left(\frac{R_g}{l_{\text{cycle}}}\right)^2 \tag{S35A}$$

$$= 4RT\left(\frac{R_{\text{ee}}/6^{1/2}}{D\left(\frac{\pi}{6}\right)^{1/3}}\right)^2 \tag{S35B}$$

$$= \frac{4}{6}\left(\frac{\pi}{6}\right)^{-2/3} RT\left(\frac{R_{\text{ee}}}{D}\right)^2 \tag{S35C}$$

$$= \frac{2}{3}\left(\frac{\pi}{6}\right)^{-2/3} RT\frac{N}{g} \tag{S35D}$$

$$\approx 1.03RT\frac{N}{g} \tag{S35E}$$

where we have used $\nu = 0.5$ for consistency with [12]. Thus, when expressed in our notation, this network-based relation yields a prefactor within experimental error of 1 ($1.03 \approx 1$), justifying our calculation of effective mesh size from the $\sigma$-value.

##### 6.2 Linearity

Our experiments suggest compression blob sizes ranging from 150-2000 residues. Thus, even for our largest chain (400AA), $N/g$ takes values from 0.2-2.6 with an average of 1. To assess how much this should impact our fitted $\sigma$-values and $\Delta G^{\text{otr}}_{\text{int.}}$, we fit to the Casassa equation[8] ($\propto \ln\left(\frac{8}{\pi^2}\sum_{i=0}^{\infty}\frac{1}{(2i+1)^2}e^{-(2i+1)^2x}\right)$, where $x = \frac{\sigma N}{RT}$ and the series is truncated at nine terms) representing the full non-linear description of the confinement free energy for a random walk polymer between two parallel plates shown to fit porous materials[8, 7]. Fitting to the Casassa equation yields small changes overall (Fig. SI1a) in $\sigma$-value (difference -0.22±0.02 cal/mol/AA, $R^2 > 0.998$, Fig. SI1b) and in $\Delta G^{\text{otr}}_{\text{int.}}$ (difference -0.04±0.01 kcal/mol, $R^2 > 0.999$, Fig. SI1c). The resulting effective local mesh sizes are $1.4 \pm 0.8$ nm larger. These minor differences suggest that corrections to being outside of the narrow confinement limit ($g \ll N$) do not impact the conclusions of our work and support the use of the linear approximation over the experimental range of chain lengths.

##### 6.3 Scaling law

To assess whether our inert random chains follow the empirical scaling law from [11], we compared those values from that calculated based on ALBATROSS[13]. We find that our scaling laws are essentially identical to ALBATROSS calculations with an RMS difference in the chain end-to-end distance of 0.32 nm (Fig. SI2a). Additionally, to assess the error of possible expansion or collapse of the polymer in the

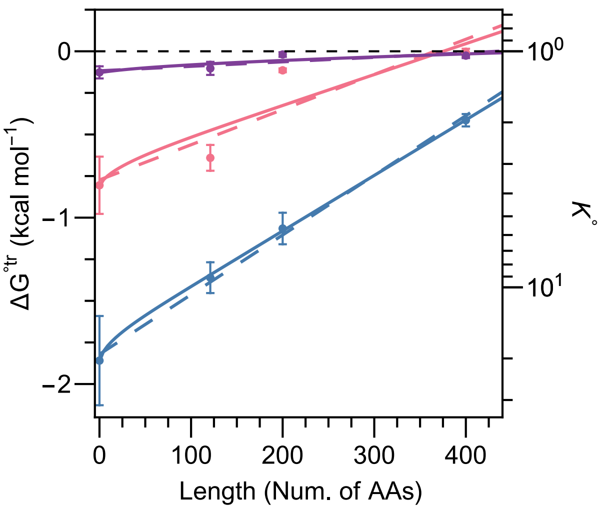

(a)

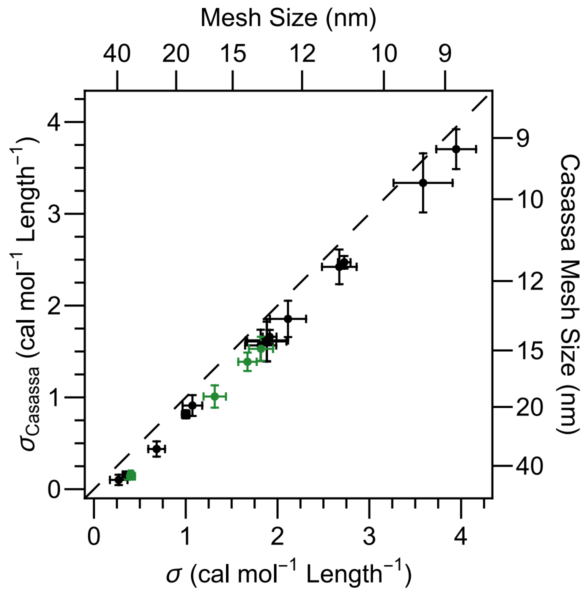

(b)

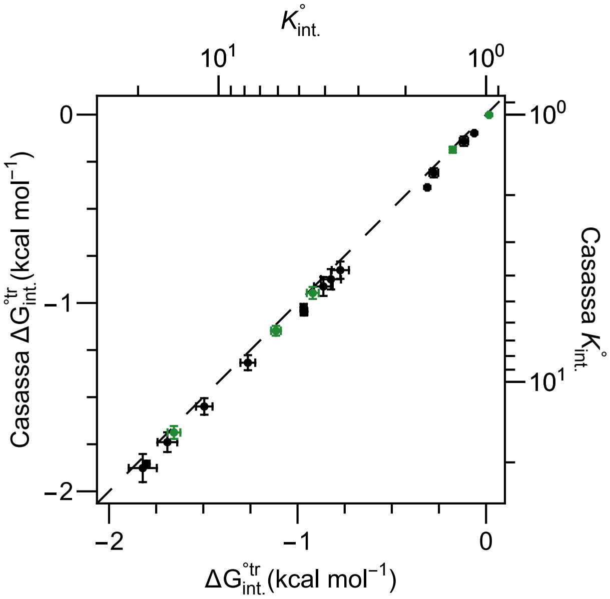

(c)

**Figure SI1: Negligible differences between non-linear and linear fit $\sigma$-values.**

(a) LSE fits using the non-linear Casassa equation (solid) and linear approximation (dashed) for proteins NGDN (blue), RPS11 (pink), and RPS3 (purple). Plot analogous to Fig. 2B(top).

(b-c) Comparison between (b)$\sigma$-values and (c) $\Delta G^{\circ tr}_{\text{int.}}$ extracted from linear and non-linear Casassa equation fits. Black points are for nucleolar proteins and green are for stress granule proteins

Figure SI1

condensate due to changes in solvent quality within the condensate compared to water, we asked about the error in mesh size anticipated based on slight expansion to $\nu$=0.57 (Fig. SI2c) and contraction to $\nu$=0.53 (Fig. SI2b) from $\nu$=0.55 representing the anticipated changes in 0.5M GdnHCl and 0.25M Sarcosine from water, respectively[11]. Employing protein-derived scaling laws[14], we calculated the effective local mesh sizes for these increases and decreases in solvent quality, finding that they result in a median change increase or decrease in mesh size of 1.0 nm and -1.3 nm, respectively. While we do not anticipate the solvent qualities to change this much, these calculations provide bounds on the accuracy of our measurements, which are on the order of our experimental error.

### 7 Elucidating microenvironment heterogeneity via $\sigma$-value and LSE

By interpreting the $\sigma$-value as a local environmental property (e.g., an effective local mesh size), we can assess heterogeneity among the microenvironments present within the condensates studied (nucleolus/GC, stress granule, and nuclear pore). Importantly, we need not invoke multiple phases to explain these differences. Phase identity is defined using endogenous compartment markers and is therefore independent of the specific probe used to measure $\sigma$. Instead, we propose that the proteins studied induce long-range correlations in the surrounding biomolecular network, thereby shaping the local biophysical properties of their immediate environment. If biological condensates behaved like simple molecular liquids, where correlations in the local environment decay within only a few molecular diameters, as quantified by pair correlation functions or radial distribution functions, we would expect $\sigma$-values measured around different proteins to converge to a single global value. However, this expectation is inconsistent with our measurements, which reveal approximately a five-fold range in $\sigma$-values surrounding different nucleolar and stress-granule proteins. This striking heterogeneity suggests that condensates are far from uniform and that individual biomolecules can actively influence the physical environment in their vicinity. We call this statistical impact of the protein on its microenvironment, microenvironmental coupling, and propose that such local structuring may have important physical–chemical implications for biomolecular behavior within condensates, including effects on assembly pathways, reaction environments, and potentially long-range regulatory mechanisms. These results are consistent with the emerging view that biomolecular condensates behave as networked fluids, in which biomolecules generate structured interaction fields that extend beyond their immediate contact partners [15, 16, 17].

### 8 Model for local energetics as a function of mesh size during ribosome biogenesis

To provide a minimal physical picture linking ribosomal intermediate maturation to the local transfer free energy, we constructed a coarse-grained energetic model based on interactions between a ribosomal intermediate and the surrounding meshwork. We decompose the local transfer free energy into contributions from three classes of interaction sites: contacts formed between the ribosomal intermediate and the local mesh, unsatisfied interaction sites remaining on the ribosomal intermediate, and unsatisfied interaction sites remaining on the mesh. Denoting these quantities by $N_{\mathrm{Cont}}$, $N_{\mathrm{RI}}$, and $N_{\mathrm{M}}$, respectively, we write

$$\Delta G^{\mathrm{tr}}(N_{\mathrm{Cont}}, N_{\mathrm{RI}}, N_{\mathrm{M}}) = \epsilon_{\mathrm{Cont}} N_{\mathrm{Cont}} + \epsilon_{\mathrm{RI}} N_{\mathrm{RI}} + \epsilon_{\mathrm{M}} N_{\mathrm{M}}. \tag{S36}$$

Within the physical picture developed in the main text, contacts between the mesh and exposed rRNA segments are favorable, whereas unsatisfied interactions are unfavorable. Accordingly, we take

$$\epsilon_{\mathrm{Cont}} < 0, \qquad \epsilon_{\mathrm{RI}} > 0, \qquad \epsilon_{\mathrm{M}} > 0. \tag{S37}$$

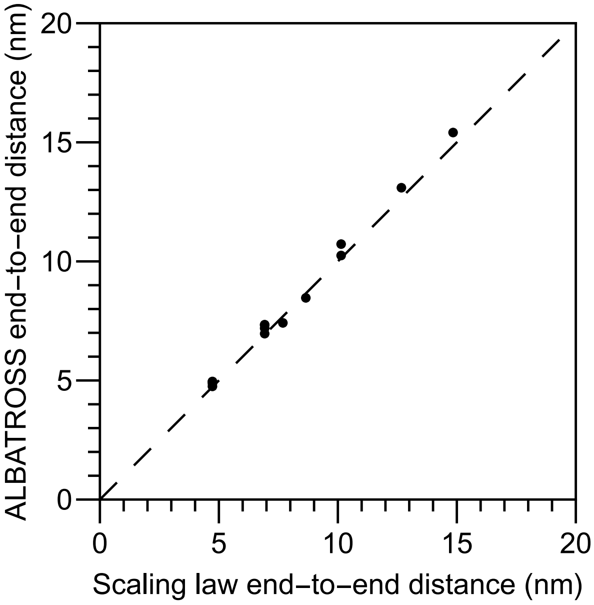

(a)

**Figure SI2: Differences in modeling chains have minimal impact on effective local mesh sizes calculated.**

(a) End-to-end distances of chains calculated using scaling laws for disordered amino-acid chains in water[11] and ALBATROSS[13].

(b-c) Comparison between extracted effective mesh sizes calculated due to an (b) increase and (c) decrease in solvent quality compared to that in water. Brown points are for nucleolar proteins, green are for stress granule proteins, and blue are for nuclear pore proteins.

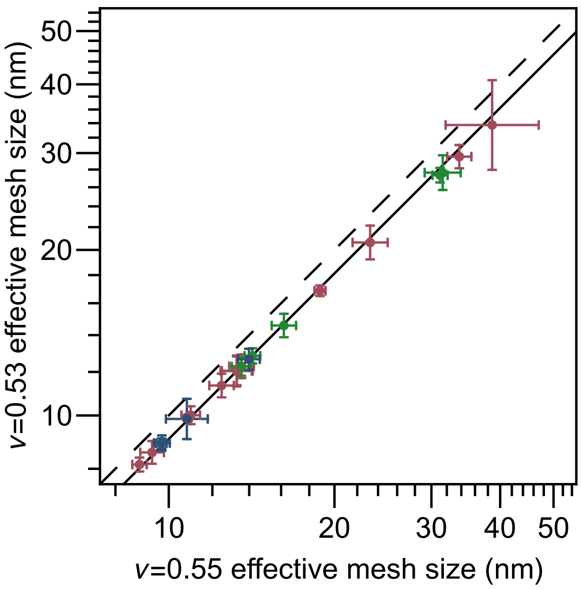

(b)

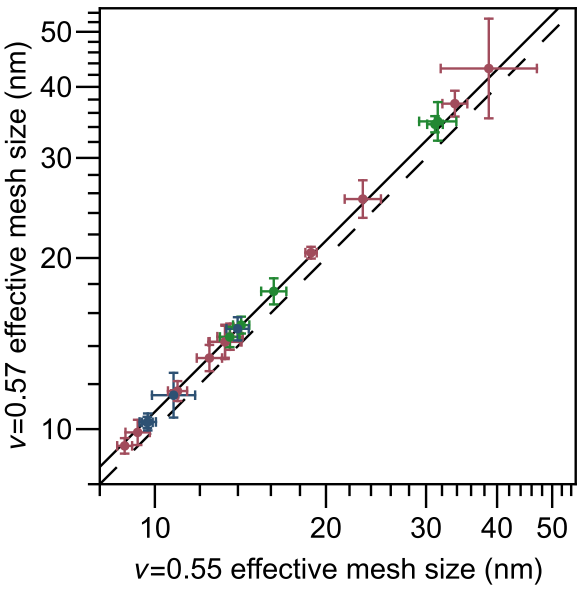

(c)

Figure SI2

We next impose conservation of interaction sites by defining the maximum number of available interaction sites on the ribosomal intermediate and the mesh as $N_{\mathrm{RI}}^{\max}$ and $N_{\mathrm{M}}^{\max}$, respectively:

$$N_{\mathrm{RI}} = N_{\mathrm{RI}}^{\max} - N_{\mathrm{Cont}}, \tag{S38A}$$

$$N_{\mathrm{M}} = N_{\mathrm{M}}^{\max} - N_{\mathrm{Cont}}. \tag{S38B}$$

This formulation reduces the model to a single independent variable, the number of formed contacts $N_{\mathrm{Cont}}$, with all other terms determined by the available valences.

To obtain a closed form, we assume that contacts are maximized locally, such that

$$N_{\mathrm{Cont}} = \min(N_{\mathrm{RI}}^{\max}, N_{\mathrm{M}}^{\max}). \tag{S39}$$

This corresponds to a saturation regime in which interactions are sufficiently favorable that all available binding capacity is utilized, limited only by the smaller of the two valences.

Substituting these relations yields

$$\Delta G^{\mathrm{tr}}(N_{\mathrm{RI}}^{\max}, N_{\mathrm{M}}^{\max}) = \epsilon_{\mathrm{RI}} N_{\mathrm{RI}}^{\max} + \epsilon_{\mathrm{M}} N_{\mathrm{M}}^{\max} + (\epsilon_{\mathrm{Cont}} - \epsilon_{\mathrm{RI}} - \epsilon_{\mathrm{M}}) \min(N_{\mathrm{RI}}^{\max}, N_{\mathrm{M}}^{\max}). \tag{S40}$$

This expression naturally separates into two regimes depending on which component limits contact formation. When $N_{\mathrm{RI}}^{\max} < N_{\mathrm{M}}^{\max}$, the ribosomal intermediate is limiting and the free energy scales with its available interaction valence. Conversely, when $N_{\mathrm{M}}^{\max} < N_{\mathrm{RI}}^{\max}$, the mesh limits contact formation and the free energy instead reflects the local mesh capacity. Thus, the model predicts a crossover between ribosomal-intermediate-limited and mesh-limited regimes.

To generate the schematic in Fig. 4C and Fig. S4, we model $N_{\mathrm{RI}}^{\max}$ as a discrete quantity reflecting the number of expanded rRNA segments available for interaction at different stages of ribosome biogenesis, using values from 0 to 3. In contrast, $N_{\mathrm{M}}^{\max}$ is treated as a continuous variable that increases with local mesh contact density. For illustration, we choose

$$\epsilon_{\mathrm{Cont}} = -RT, \qquad \epsilon_{\mathrm{M}} = RT, \qquad \epsilon_{\mathrm{RI}} = 12RT. \tag{S41}$$

These parameter values impose a strong energetic penalty for unsatisfied ribosomal-intermediate interaction sites relative to the mesh, biasing the system toward maximizing contacts with the local environment.

These parameter values are not intended as a quantitative fit, but rather as a minimal phenomenological model that captures the qualitative idea that early ribosomal intermediates, which present more exposed rRNA interaction sites, stabilize tighter local mesh environments, whereas later intermediates progressively reduce this valence and thereby weaken local mesh stabilization.

### 9 Domain decomposition and nearest-neighbor coupling

The statistical mechanical framework above can be extended to make explicit how local coupling between neighboring elements contributes to insertion free energies. Consider two elements of a fused probe, $\alpha$ and $\beta$, inserted jointly into phase $i$. The corresponding excess free energy is

$$G_{\alpha-\beta}^{\mathrm{ex},i} = -RT \ln \langle w(\alpha,\beta) \rangle_0^i, \tag{S42}$$

where

$$\langle w(\alpha,\beta) \rangle_0^i = \int P^i(\Omega)\, w^i(\alpha,\beta|\Omega)\, d\Omega \tag{S43}$$

is the average insertion weight of the composite probe.

To compare this with the insertion of $\alpha$ and $\beta$ separately, we define the nearest-neighbor coupling free energy

$$\boxed{\begin{aligned} G^{\mathrm{ex},i}_{\mathrm{NN},\alpha-\beta} &\equiv G^{\mathrm{ex},i}_{\alpha-\beta} - G^{\mathrm{ex},i}_{\alpha} - G^{\mathrm{ex},i}_{\beta} \qquad \text{(S44A)} \\ &= -RT \ln \frac{\langle w(\alpha,\beta)\rangle_0^i}{\langle w(\alpha)\rangle_0^i \, \langle w(\beta)\rangle_0^i}. \qquad \text{(S44B)} \end{aligned}}$$

When $G^{\mathrm{ex},i}_{\mathrm{NN},\alpha,\beta} = 0$, the insertion free-energy cost is additive. Deviations from zero, therefore, quantify non-additive coupling between neighboring elements of the probe.

To interpret this quantity, it is useful to introduce

$$\langle w(\alpha) w(\beta)\rangle_0^i = \int P^i(\Omega)\, w^i(\alpha|\Omega)\, w^i(\beta|\Omega)\, d\Omega. \tag{S45}$$

This allows the coupling free energy to be decomposed as

$$G^{\mathrm{ex},i}_{\mathrm{NN},\alpha,\beta} = -RT \ln \left( \frac{\langle w(\alpha,\beta)\rangle_0^i}{\langle w(\alpha) w(\beta)\rangle_0^i} \right) - RT \ln \left( \frac{\langle w(\alpha) w(\beta)\rangle_0^i}{\langle w(\alpha)\rangle_0^i \, \langle w(\beta)\rangle_0^i} \right). \tag{S46}$$

The first term,

$$-RT \ln \left( \frac{\langle w(\alpha,\beta)\rangle_0^i}{\langle w(\alpha) w(\beta)\rangle_0^i} \right), \tag{S47}$$

measures the extent to which the joint insertion weight fails to factorize within a fixed background microstate $\Omega$. It therefore reports an explicit *residual coupling* between $\alpha$ and $\beta$, such as steric interference or direct local interactions between neighboring elements.

The second term,

$$-RT \ln \left( \frac{\langle w(\alpha) w(\beta)\rangle_0^i}{\langle w(\alpha)\rangle_0^i \, \langle w(\beta)\rangle_0^i} \right), \tag{S48}$$

has a probabilistic interpretation. It measures whether microstates favorable for inserting $\alpha$ also tend to be favorable for inserting $\beta$. Equivalently, it quantifies a *correlational coupling* between the random variables $w(\alpha|\Omega)$ and $w(\beta|\Omega)$ over the ensemble of background microstates. Thus, even if

$$w^i(\alpha, \beta|\Omega) \approx w^i(\alpha|\Omega)\, w^i(\beta|\Omega), \tag{S49}$$

so that the residual coupling is negligible, the total coupling free energy can still be nonzero if $\alpha$ and $\beta$ are correlated across microenvironments.

This distinction is useful physically. Residual coupling reflects direct or short-range interactions between neighboring elements within a given local environment, whereas correlational coupling reflects shared preference (or resistance) for the same classes of condensate microenvironments. In the context of the $\sigma$-value measurements used here, the $\sigma$-value integrates both residual and correlational contributions, although we expect the dominant contribution to arise from correlational coupling. In this view, $\sigma_\alpha$ reports how conditioning on the anchor $\alpha$ reweights the ensemble of condensate microenvironments, and whether the resulting microenvironments are more or less accommodating to an appended residue.

In fact, the nearest-neighbor coupling can be simplified in terms of $\sigma$-values by letting $\beta$ be a chain of $N$ residues and expanding to first order in chain length $N$. In this case, the nearest-neighbor coupling free energy becomes

$$G^{\text{ex},i}_{\text{NN},\alpha,\beta} \equiv G^{\text{ex},i}_{\alpha-\beta} - G^{\text{ex},i}_{\alpha} - G^{\text{ex},i}_{\beta} \tag{S50A}$$

$$= -RT \ln \frac{\langle w(\beta)\rangle^i_\alpha}{\langle w(\beta)\rangle^i_0} \tag{S50B}$$

$$\approx (\sigma_\alpha - \sigma_0)N \tag{S50C}$$

where we have employed equation S29.

Thus, $(\sigma_\alpha - \sigma_0)$ defines the change in the marginal free energy cost of inserting an additional residue upon conditioning on $\alpha$. In the case of a chain primarily sensitive to confinement, as considered here, this quantity reports the difference between the effective local confinement around $\alpha$ and that of the average condensate environment. The sign of this difference determines whether the microenvironments favored by $\alpha$ are more or less accommodating to the appended chain.

#### 10 Theoretical basis for Length-Dependent Spatial Coupling

The statistical framework developed above can be extended beyond nearest-neighbor coupling to include next-nearest-neighbor coupling, which we term spatial coupling. Notably, we will relate the spatial coupling to the pair correlation function, g(r), which describes the spatial enrichment between molecules as a function of distance and the corresponding energetic analog, the potential of mean force. Together, these relationships provide a partitioning based framework to interrogate spatial organization between proteins. Below, we begin by briefly introducing those concepts.

The pair correlation function can be defined in the context of the equations herein, as a ratio of the average insertion weight (equation S16) of two components $\alpha$ and $\beta$ jointly inserted at a fixed distance $r$, denoted $\langle w(\alpha, r, \beta)\rangle^i_0$, compared to the independent insertion weights,

$$\boxed{g^i_{\alpha,\beta}(r) = \frac{\langle w(\alpha, r, \beta)\rangle^i_0}{\langle w(\alpha)\rangle^i_0 \langle w(\beta)\rangle^i_0}} \tag{S51}$$

and where

$$G^i_{\alpha,\beta}(r) = -RT \ln(g^i_{\alpha,\beta}(r)) \tag{S52A}$$

$$= -RT \ln \left( \frac{\langle w(\alpha, r, \beta)\rangle^i_0}{\langle w(\alpha)\rangle^i_0 \langle w(\beta)\rangle^i_0} \right) \tag{S52B}$$

is referred to as the potential of mean force, describing the mean potential at a separation, r, relative to that at infinite separation[18]. This can be seen by evaluating that limit:

$$\lim_{r\to\infty} g^i_{\alpha,\beta}(r) = \lim_{r\to\infty} \frac{\langle w(\alpha, r, \beta)\rangle^i_0}{\langle w(\alpha)\rangle^i_0 \langle w(\beta)\rangle^i_0} \tag{S53A}$$

$$= \frac{\langle w(\alpha, \infty, \beta)\rangle^i_0}{\langle w(\alpha)\rangle^i_0 \langle w(\beta)\rangle^i_0} \tag{S53B}$$

$$= \frac{\langle w(\alpha)\rangle^i_0 \langle w(\beta)\rangle^i_0}{\langle w(\alpha)\rangle^i_0 \langle w(\beta)\rangle^i_0} \tag{S53C}$$

$$= 1 \tag{S53D}$$

where we have employed the relationship, $\langle w(\alpha,\infty,\beta)\rangle_0^i = \langle w(\alpha)\rangle_0^i \langle w(\beta)\rangle_0^i$, signifying that at infinite distances, the weight of insertion between $\alpha$ and $\beta$ will become uncorrelated. Indeed, plugging equation S53D into equation S52B implies that the potential of mean force at infinite separation is zero. Finally, we remind the reader that the $g_{\alpha,\beta}(r)$ is not abstract; it describes the precise spatial enrichment between $\alpha$ and $\beta$ as a function of distance. Now with this in mind, we return to the coupling free energy attempting to relate it to $g_{\alpha,\beta}(r)$.

We begin by evaluating the coupling free energy that remains from the composite probe $\alpha$-$L$-$\beta$ after intrinsic domain and nearest-neighbor domain couplings are subtracted. Proceeding:

$$G^{ex}_{Sp.Co.} = G^{ex}_{\alpha-L-\beta} - G^{ex}_{\alpha} - G^{ex}_{L} - G^{ex}_{\beta} - G^{ex}_{NN,\alpha-L} - G^{ex}_{NN,L-\beta} \tag{S54A}$$

$$= G^{ex}_{\alpha-L-\beta} - G^{ex}_{\alpha} - G^{ex}_{L} - G^{ex}_{\beta} - (G^{ex}_{\alpha-L} - G^{ex}_{\alpha} - G^{ex}_{L}) - (G^{ex}_{L-\beta} - G^{ex}_{L} - G^{ex}_{\beta}) \tag{S54B}$$

$$= G^{ex}_{\alpha-L-\beta} - G^{ex}_{\alpha-L} - G^{ex}_{L-\beta} + G^{ex}_{L} \tag{S54C}$$

where we have employed equation S44A. Using the sequential insertion form introduced above (Eq. S21B), this can be further reduced to:

$$G^{ex}_{Sp.Co.} = (G^{ex}_{L} - RT\ln\langle w(\alpha,\beta)\rangle_L^i) - (G^{ex}_{L} - RT\ln\langle w(\alpha)\rangle_L^i) - (G^{ex}_{L} - RT\ln\langle w(\beta)\rangle_L^i) + G^{ex}_{L} \tag{S54D}$$

$$= -RT\ln\left(\frac{\langle w(\alpha,\beta)\rangle_L^i}{\langle w(\alpha)\rangle_L^i \langle w(\beta)\rangle_L^i}\right) \tag{S54E}$$

$$= -RT\ln\left(g^i_{\alpha,\beta}(L)\right) \tag{S54F}$$

and where

$$\boxed{g^i_{\alpha,\beta}(L) = \frac{\langle w(\alpha,\beta)\rangle_L^i}{\langle w(\alpha)\rangle_L^i \langle w(\beta)\rangle_L^i}} \tag{S55}$$

is defined due to its notable similarity with $g^i_{\alpha,\beta}(r)$ defined in equation S51 .

$G^{ex}_{Sp.Co.}$ and $g^i_{\alpha,\beta}(L)$ therefore quantify the thermodynamic energy and enrichment due to the coupling between $\alpha$ and $\beta$ when connected by a domain "$L$". This intuitively makes sense, as the notation $\langle w(...)\rangle_L^i$ is the average Boltzmann weight of adding other components ("...") into a micro-state that already contains $L$. In our case, where $L$ is a disordered chain of N residues (referred to as the linker), then $g^i_{\alpha,\beta}(L)$ will have a limit similar at large chain lengths to that of $g^i_{\alpha,\beta}(r)$ at large distances. Expanding similarly to equation S53D:

$$\lim_{N\to\infty} g^i_{\alpha,\beta}(L) = \lim_{N\to\infty} \frac{\langle w(\alpha,\beta)\rangle_L^i}{\langle w(\alpha)\rangle_L^i \langle w(\beta)\rangle_L^i} \tag{S56A}$$

$$\approx \lim_{N\to\infty} \frac{\langle w(\alpha)w(\beta)\rangle_L^i}{\langle w(\alpha)\rangle_L^i \langle w(\beta)\rangle_L^i} \tag{S56B}$$

$$= \frac{\langle w(\alpha)\rangle_L^i \langle w(\beta)\rangle_L^i}{\langle w(\alpha)\rangle_L^i \langle w(\beta)\rangle_L^i} \tag{S56C}$$

$$= 1 \tag{S56D}$$

where we assume that the linker results in a near zero residual coupling (as above, see equation S47) and have employed the relationship, $\lim_{N\to\infty} \langle w(\alpha)w(\beta)\rangle_L^i = \langle w(\alpha)\rangle_L^i \langle w(\beta)\rangle_L^i$, as at infinite linker lengths, the weight of insertion between $\alpha$ and $\beta$ will become uncorrelated.

Thus, $g^i_{\alpha,\beta}(L)$ can be related to $g^i_{\alpha,\beta}(r)$ under two additional approximations: (1) the linker samples separations between $\alpha$ and $\beta$ according to its bare end-to-end distance distribution $P^0(r|L)$, and (2) the spatial correlations between $\alpha$ and $\beta$ at fixed separation $r$ are intrinsic properties of the condensate microenvironment, independent of the particular linker used to probe them. Under these assumptions, the joint insertion weight for $\alpha$ and $\beta$ connected by linker $L$ can be written as

$$\langle w(\alpha, \beta)\rangle^i_L \approx \langle w(\alpha)\rangle^i_L \, \langle w(\beta)\rangle^i_L \int P^0(r|L)\, g^i_{\alpha,\beta}(r)\, dr. \tag{S57}$$

where the factorization $\langle w(\alpha)\rangle^i_L \, \langle w(\beta)\rangle^i_L$ is exact in the limit $N \to \infty$ (equation S56D). This expression can be interpreted as averaging the distance-dependent correlations over the separations sampled by the linker, analogous to how the potential of mean force encodes distance-resolved interactions. Substituting into the definition of $g^i_{\alpha,\beta}(L)$, the marginal insertion weights cancel exactly, yielding the central result:

$$\boxed{g^i_{\alpha,\beta}(L) \approx \int P^0(r|L)\, g^i_{\alpha,\beta}(r)\, dr} \tag{S58}$$

Thus, the experimentally measured $g^i_{\alpha,\beta}(L)$ is a weighted average of the true distance-dependent pair correlation function $g^i_{\alpha,\beta}(r)$, smeared by the end-to-end distance distribution of the linker $P^0(r|L)$. In this way, LDSC provides a direct measurement of correlational microenvironment compatibility between $\alpha$ and $\beta$ as a function of their spatial separation, with the linker setting the length scale over which the condensate is probed.

Notably, $g^i_{\alpha,\beta}(L)$ as derived here measures predominantly *correlational coupling* between $\alpha$ and $\beta$ rather than residual coupling arising from direct interactions between them. This follows directly from the cancellation of the single-element insertion weights in the derivation above, which absorbs all direct insertion energetics into the reference terms, leaving only the relational quantity $g^i_{\alpha,\beta}(r)$. In this sense, LDSC is the spatial analog of the correlational coupling defined in equation S48, extended to quantify how microenvironment compatibility varies as a function of the distance between two proteins.

In summary, the behavior of $g^i_{\alpha,\beta}(L)$ reflects how the condensate organizes different components in space. The function $g^i_{\alpha,\beta}(r)$ encodes whether $\alpha$ and $\beta$ preferentially occur near one another ($g > 1$) or avoid one another ($g < 1$) at a given separation $r$. Because the linker samples a distribution of separations set by its length, $g^i_{\alpha,\beta}(L)$ reports the average of these spatial preferences over the distances that the linker can access. When the linker is short, it primarily samples small separations, and $g^i_{\alpha,\beta}(L)$ reflects local organization—whether $\alpha$ and $\beta$ tend to co-localize or exclude one another in their immediate surroundings. As the linker length increases, it samples progressively larger separations, and these local preferences are averaged over a broader spatial range. At sufficiently large linker lengths, any spatial organization is washed out, and $g^i_{\alpha,\beta}(L)$ approaches unity, indicating no net preference. In this way, varying the linker length provides a direct means of probing how spatial preferences between $\alpha$ and $\beta$ evolve with distance. Rather than measuring a single interaction, LDSC captures how the condensate organizes components across length scales, revealing whether interactions are localized or persist over extended spatial ranges.

### 11 Implementation of the linear decomposition

To quantify the energetic contributions of NPM1 domains and related constructs used in the LDSC analysis, we employ a linear decomposition of the transfer free energy. Notably, we focus on the 3-domain structure which we will abbreviate "N" - "L" - "C" as done in the schematics (Fig. 3, S6). We decompose the transfer free energy into three classes of terms: intrinsic (intr.) domain contributions, nearest-neighbor coupling (NN.Co.) terms, and a next-nearest-neighbor term that we refer to as the spatial coupling (Sp.Co.).

Specifically, we write the transfer free energies of the measured constructs as

$$\Delta G^{\text{otr}}_{\text{FP}} = \Delta G^{\text{otr}}_{\text{intr.FP}} \tag{S59A}$$
$$\Delta G^{\text{otr}}_{\text{L}} = \Delta G^{\text{otr}}_{\text{intr.L}} + \Delta G^{\text{otr}}_{\text{intr.FP}} \tag{S59B}$$
$$\Delta G^{\text{otr}}_{\text{N}} = \Delta G^{\text{otr}}_{\text{intr.N}} + \Delta G^{\text{otr}}_{\text{intr.FP}} \tag{S59C}$$
$$\Delta G^{\text{otr}}_{\text{C}} = \Delta G^{\text{otr}}_{\text{intr.C}} + \Delta G^{\text{otr}}_{\text{intr.FP}} \tag{S59D}$$
$$\Delta G^{\text{otr}}_{\text{NL}} = \Delta G^{\text{otr}}_{\text{intr.N}} + \Delta G^{\text{otr}}_{\text{intr.L}} + \Delta G^{\text{otr}}_{\text{intr.FP}} + \Delta G^{\text{otr}}_{\text{NN.Co.,N\&L}} \tag{S59E}$$
$$\Delta G^{\text{otr}}_{\text{LC}} = \Delta G^{\text{otr}}_{\text{intr.L}} + \Delta G^{\text{otr}}_{\text{intr.C}} + \Delta G^{\text{otr}}_{\text{intr.FP}} + \Delta G^{\text{otr}}_{\text{NN.Co.,L\&C}} \tag{S59F}$$
$$\begin{aligned}\Delta G^{\text{otr}}_{\text{NLC}} = {} & \Delta G^{\text{otr}}_{\text{intr.N}} + \Delta G^{\text{otr}}_{\text{intr.L}} + \Delta G^{\text{otr}}_{\text{intr.C}} + \Delta G^{\text{otr}}_{\text{intr.FP}} \\ & + \Delta G^{\text{otr}}_{\text{NN.Co.,N\&L}} + \Delta G^{\text{otr}}_{\text{NN.Co.,L\&C}} + \Delta G^{\text{otr}}_{\text{Sp.Co.}}.\end{aligned} \tag{S59G}$$

These relations allow the intrinsic and coupling terms to be solved directly from the measured construct energies:

$$\Delta G^{\text{otr}}_{\text{intr.FP}} = \Delta G^{\text{otr}}_{\text{FP}} \tag{S60A}$$
$$\Delta G^{\text{otr}}_{\text{intr.C}} = \Delta G^{\text{otr}}_{\text{C}} - \Delta G^{\text{otr}}_{\text{FP}} \tag{S60B}$$
$$\Delta G^{\text{otr}}_{\text{intr.N}} = \Delta G^{\text{otr}}_{\text{N}} - \Delta G^{\text{otr}}_{\text{FP}} \tag{S60C}$$
$$\Delta G^{\text{otr}}_{\text{intr.L}} = \Delta G^{\text{otr}}_{\text{L}} - \Delta G^{\text{otr}}_{\text{FP}} \tag{S60D}$$
$$\Delta G^{\text{otr}}_{\text{NN.Co.,N\&L}} = \Delta G^{\text{otr}}_{\text{NL}} - \Delta G^{\text{otr}}_{\text{N}} - \Delta G^{\text{otr}}_{\text{L}} + \Delta G^{\text{otr}}_{\text{FP}} \tag{S60E}$$
$$\Delta G^{\text{otr}}_{\text{NN.Co.,L\&C}} = \Delta G^{\text{otr}}_{\text{LC}} - \Delta G^{\text{otr}}_{\text{L}} - \Delta G^{\text{otr}}_{\text{C}} + \Delta G^{\text{otr}}_{\text{FP}} \tag{S60F}$$
$$\Delta G^{\text{otr}}_{\text{Sp.Co.}} = \Delta G^{\text{otr}}_{\text{NLC}} - \Delta G^{\text{otr}}_{\text{NL}} - \Delta G^{\text{otr}}_{\text{LC}} + \Delta G^{\text{otr}}_{\text{L}}. \tag{S60G}$$

Errors on the derived quantities were propagated by summing the variances of the contributing terms. For experimental practicality, we neglect couplings between the fluorescent protein and other domains. Importantly, such terms cancel in the expression for the spatial coupling, rendering the LDSC measurement maximally robust to complexities introduced by the fluorescent tag.

This stepwise representation is analogous to standard decompositions of free energies into intrinsic and pairwise interaction terms, where non-additive contributions are captured as explicit coupling terms (see [19]). For schematic presentation in Fig. 3A, the same decomposition may be written in a simplified stepwise form:

$$\Delta G^{\text{otr}}_{\text{N}} = \Delta G^{\text{otr}}_{\text{FP}} + \Delta G^{\text{otr}}_{\text{intr.N}} \tag{S61A}$$
$$\Delta G^{\text{otr}}_{\text{L}} = \Delta G^{\text{otr}}_{\text{FP}} + \Delta G^{\text{otr}}_{\text{intr.L}} \tag{S61B}$$
$$\Delta G^{\text{otr}}_{\text{C}} = \Delta G^{\text{otr}}_{\text{FP}} + \Delta G^{\text{otr}}_{\text{intr.C}} \tag{S61C}$$
$$\Delta G^{\text{otr}}_{\text{NL}} = \Delta G^{\text{otr}}_{\text{L}} + \Delta G^{\text{otr}}_{\text{intr.N}} + \Delta G^{\text{otr}}_{\text{NN.Co.,N\&L}} \tag{S61D}$$
$$\Delta G^{\text{otr}}_{\text{LC}} = \Delta G^{\text{otr}}_{\text{L}} + \Delta G^{\text{otr}}_{\text{intr.C}} + \Delta G^{\text{otr}}_{\text{NN.Co.,L\&C}} \tag{S61E}$$
$$\Delta G^{\text{otr}}_{\text{NLC}} = \Delta G^{\text{otr}}_{\text{NL}} + \Delta G^{\text{otr}}_{\text{intr.C}} + \Delta G^{\text{otr}}_{\text{NN.Co.,L\&C}} + \Delta G^{\text{otr}}_{\text{Sp.Co.}}. \tag{S61F}$$

This decomposition provides a compact way to separate the partitioning contributions of individual domains from the non-additive effects that arise when they are combined within the same molecule.

#### 12 Theoretical basis for studying small condensates

Many condensates in cells are diffraction limited. To understand how the diffraction limit impacts the ability to extract transfer free energies, we sought a simple theoretical framework. For simplicity, we assume that the signal arises from two underlying phases, the dilute and dense phases, which are not spatially resolved. Because the condensate is diffraction limited, measurements of the dilute phase, $C_{\text{dil}}$, are taken to represent the true dilute phase concentration. In contrast, only a fraction $f_{\text{den}}$ of the dense-phase measurement arises from the dense phase itself, while the remaining fraction $1 - f_{\text{den}}$ reflects signal originating from the surrounding dilute phase due to point-spread-function mixing. Thus, if $C_{\text{den}}^{\text{true}}$ denotes the true, super-resolved, dense-phase concentration, the measured dense-phase concentration is

$$C_{\text{den}} = f_{\text{den}} C_{\text{den}}^{\text{true}} + (1 - f_{\text{den}}) C_{\text{dil}}. \tag{S62}$$

The apparent transfer free energy between dilute and dense phases is therefore

$$\begin{aligned} \Delta G^{\text{tr,app}} &= -RT \ln \left( \lim_{C_{\text{dil}} \to 0} \frac{C_{\text{den}}}{C_{\text{dil}}} \right) && \text{(S63A)} \\ &= -RT \ln \left( \lim_{C_{\text{dil}} \to 0} \frac{f_{\text{den}} C_{\text{den}}^{\text{true}} + (1 - f_{\text{den}}) C_{\text{dil}}}{C_{\text{dil}}} \right) && \text{(S63B)} \\ &= -RT \ln \left( f_{\text{den}} \left( \lim_{C_{\text{dil}} \to 0} \frac{C_{\text{den}}^{\text{true}}}{C_{\text{dil}}} \right) + (1 - f_{\text{den}}) \right) && \text{(S63C)} \\ &= -RT \ln \left( f_{\text{den}} K^{\circ\text{tr}} + (1 - f_{\text{den}}) \right), && \text{(S63D)} \end{aligned}$$

where $K^{\circ\text{tr}}$ is the true transfer equilibrium constant defined in the absence of diffraction-limited mixing. This expression highlights two limiting regimes:

$$\boxed{\begin{aligned} \lim_{K^{\circ\text{tr}} \gg 1} \Delta G^{\text{tr,app}} &= -RT \ln \left( f_{\text{den}} K^{\circ\text{tr}} \right), && \text{(S64A)} \\ \lim_{K^{\circ\text{tr}} \ll 1} \Delta G^{\text{tr,app}} &= -RT \ln \left( 1 - f_{\text{den}} \right). && \text{(S64B)} \end{aligned}}$$

Thus, in the former, high partitioning molecules yield a transfer free energy that is offset by $-RT \ln (f_{\text{den}})$ and in the latter, the apparent transfer free energy saturates at a finite value set by the diffraction-limited mixing fraction $-RT \ln (1 - f_{\text{den}})$, even when the true partitioning becomes arbitrarily unfavorable. More generally, diffraction-limited measurements report a log-sum of contributions from spatially distinct sources rather than a single thermodynamic phase, as reflected in the logarithm of a weighted sum in the expression above. This form reflects the fact that measured concentrations are linear mixtures of underlying phases, whereas the inferred transfer free energy depends logarithmically on their ratio, leading to systematic compression of strong partitioning and saturation of weak partitioning.

### 13 Diffraction-limited model for nuclear pore measurements

Following the diffraction-limited mixing framework above, we treat the measured nuclear pore signal as a weighted sum of unresolved contributions from the pore, nucleoplasm (np), and cytoplasm (cp). Thus, the measured signal does not arise from a single spatially resolved phase. Instead, the apparent pore concentration is written as

$$C_{\text{pore}}^{\text{app}} = f_{\text{pore}} C_{\text{pore}}^{\text{true}} + f_{\text{np}} C_{\text{np}} + f_{\text{cp}} C_{\text{cp}}, \tag{S65}$$

where $f_{\text{pore}} + f_{\text{np}} + f_{\text{cp}} = 1$. These fractions represent the relative contributions of each compartment to the measured pore-region intensity due to point-spread-function mixing.

We define the apparent transfer free energy of the nuclear pore relative to the cytoplasm as

$$\Delta G^{\text{otr,app}}_{\text{pore}\leftarrow\text{cp}} = -RT \ln \left( \lim_{C_{\text{cp}}\to 0} \frac{C^{\text{app}}_{\text{pore}}}{C_{\text{cp}}} \right). \tag{S66}$$

Substituting gives

$$\Delta G^{\text{otr,app}}_{\text{pore}\leftarrow\text{cp}} = -RT \ln \left( f_{\text{pore}} \frac{C^{\text{true}}_{\text{pore}}}{C_{\text{cp}}} + f_{\text{np}} \frac{C_{\text{np}}}{C_{\text{cp}}} + f_{\text{cp}} \right). \tag{S67}$$

Using the relation

$$\frac{C_a}{C_b} = e^{-\frac{\Delta G^{\text{otr}}_{a\leftarrow b}}{RT}}, \tag{S68}$$

we rewrite this expression as

$$\Delta G^{\text{otr,app}}_{\text{pore}\leftarrow\text{cp}} = -RT \ln \left( f_{\text{pore}} e^{-\frac{\Delta G^{\text{otr}}_{\text{pore}\leftarrow\text{cp}}}{RT}} + f_{\text{np}} e^{-\frac{\Delta G^{\text{otr}}_{\text{np}\leftarrow\text{cp}}}{RT}} + f_{\text{cp}} \right). \tag{S69}$$

For fitting across constructs, we index nuclear pore-associated proteins by $x$. To describe the dependence on chain length $N$, we use the same linear form employed throughout:

$$\Delta G^{\text{otr}}_{\text{pore}\leftarrow\text{cp},x}(N) = \Delta G^{\text{otr}}_{\text{pore}\leftarrow\text{cp},x} + N\sigma_{\text{pore}\leftarrow\text{cp},x}, \tag{S70}$$

$$\Delta G^{\text{otr}}_{\text{np}\leftarrow\text{cp},x}(N) = \Delta G^{\text{otr}}_{\text{np}\leftarrow\text{cp},x} + N\sigma_{\text{np}\leftarrow\text{cp},x} + \Delta\Delta G^{\text{otr}}_{x}\,\delta_{N,200}. \tag{S71}$$

Here, $\Delta G^{\text{otr}}_{\text{pore}\leftarrow\text{cp},x}$ and $\Delta G^{\text{otr}}_{\text{np}\leftarrow\text{cp},x}$ are the extrapolated transfer free energies for construct $x$ from cytoplasm to the true pore and nucleoplasm compartments, respectively, before diffraction-limited mixing. Likewise, $\sigma_{\text{pore}\leftarrow\text{cp},x}$ and $\sigma_{\text{np}\leftarrow\text{cp},x}$ describe the chain-length dependence of those underlying transfer free energies. The term $\Delta\Delta G^{\text{otr}}_{x}\,\delta_{N,200}$ allows a construct-specific offset for the $N = 200$ chain-length measurement, where $\delta_{N,200} = 1$ for $N = 200$ and 0 otherwise.

Substituting these linear forms into Eq. S69 yields the fitting form

$$\begin{aligned}\Delta G^{\text{otr,app}}_{\text{pore}\leftarrow\text{cp},x}(N) = -RT \ln \Bigg[ & f_{\text{pore}} \exp\left( -\frac{\Delta G^{\text{otr}}_{\text{pore}\leftarrow\text{cp},x} + N\sigma_{\text{pore}\leftarrow\text{cp},x}}{RT} \right) \\ & + f_{\text{np}} \exp\left( -\frac{\Delta G^{\text{otr}}_{\text{np}\leftarrow\text{cp},x} + N\sigma_{\text{np}\leftarrow\text{cp},x} + \Delta\Delta G^{\text{otr}}_{x}\,\delta_{N,200}}{RT} \right) + f_{\text{cp}} \Bigg].\end{aligned} \tag{S72}$$

In parallel, the nucleoplasm measurements are fitted directly as

$$\Delta G^{\text{otr}}_{\text{np}\leftarrow\text{cp},x}(N) = \Delta G^{\text{otr}}_{\text{np}\leftarrow\text{cp},x} + N\sigma_{\text{np}\leftarrow\text{cp},x} + \Delta\Delta G^{\text{otr}}_{x}\,\delta_{N,200}. \tag{S73}$$

The fractional contributions are parameterized to enforce normalization:

$$f_{\text{pore}} = \frac{e^{\lambda_{\text{pore}}}}{1+e^{\lambda_{\text{pore}}}}, \quad f_{\text{np}} = (1-f_{\text{pore}})\frac{1}{1+e^{\lambda_{\text{cp}}}}, \quad f_{\text{cp}} = (1-f_{\text{pore}})\frac{e^{\lambda_{\text{cp}}}}{1+e^{\lambda_{\text{cp}}}}. \tag{S74}$$

The pore-region and nucleoplasm measurements are fit jointly. The nucleoplasm measurements constrain $\Delta G^{\text{otr}}_{\text{np}\leftarrow\text{cp},x}$, $\sigma_{\text{np}\leftarrow\text{cp},x}$, and $\Delta\Delta G^{\text{otr}}_{x}$, while the apparent pore measurements constrain $\Delta G^{\text{otr}}_{\text{pore}\leftarrow\text{cp},x}$, $\sigma_{\text{pore}\leftarrow\text{cp},x}$, and the fractional signal contributions through $\lambda_{\text{pore}}$ and $\lambda_{\text{cp}}$. The fractional contributions were fit globally across all constructs, rather than separately for each construct. Thus, a single pair of parameters, $\lambda_{\text{pore}}$ and $\lambda_{\text{cp}}$, determines $f_{\text{pore}}$, $f_{\text{np}}$, and $f_{\text{cp}}$ for the full dataset. The pore-region and nucleoplasm measurements were fit jointly by nonlinear least squares using the Levenberg–Marquardt algorithm.

The fitted dataset contained both nucleoplasm-to-cytoplasm measurements and apparent pore-to-cytoplasm measurements for each construct and chain length. The linker-only construct was included as an additional construct in the joint fit. For this construct, the chain-length dependence was constrained to be shared between the pore and nucleoplasm terms, such that $\sigma_{\text{pore}\leftarrow\text{cp,Linker}} = \sigma_{\text{np}\leftarrow\text{cp,Linker}}$. The intercept terms for the linker-only construct were still fit separately for the pore and nucleoplasm contributions. A single variance estimator was used for all points, given by the median of the squared experimental uncertainties. Parameter uncertainties were obtained from the nonlinear model fit.

Thus, the apparent pore transfer free energy is a log-sum of underlying transfer free energies from spatially unresolved compartments. In the limit $f_{\text{pore}} \to 1$, the true $\Delta G^{\text{otr}}_{\text{pore}\leftarrow\text{cp}}$ is recovered, whereas for finite mixing the apparent value is systematically biased toward weaker partitioning. More generally, this model accounts for the fact that the measured pore signal is a linear mixture of compartmental signals, whereas the inferred transfer free energy depends logarithmically on the measured concentration ratio.
